# Calculation of Protein Folding Thermodynamics using Molecular Dynamics Simulations

**DOI:** 10.1101/2023.01.21.525008

**Authors:** Juan José Galano-Frutos, Francho Nerín-Fonz, Javier Sancho

**Affiliations:** Department of Biochemistry, Molecular and Cell Biology. Faculty of Science, University of Zaragoza, 50009 Zaragoza, Spain; Biocomputation and Complex Systems Physics Institute (BIFI). Joint Units BIFI-IQFR (CSIC) and GBs-CSIC, University of Zaragoza, 50018 Zaragoza, Spain; Aragon Health Research Institute (IIS Aragón), 50009 Zaragoza, Spain

## Abstract

Despite impressive advances by AlphaFold2 in the field of computational biology, the protein folding problem remains an enigma to be solved. The continuous development of algorithms and methods to explore longer simulation timescales of biological systems, as well as the enhanced accuracy of potential functions (force fields and solvent models) have not yet led to significant progress in the calculation of the thermodynamics quantities associated to protein folding from first principles. Progress in this direction can help boost related fields such as protein engineering, drug design, or genetic interpretation, but the task seems not to have been addressed by the scientific community. Following an initial explorative study, we extend here the application of a Molecular Dynamics-based approach −with the most accurate force field/water model combination previously found (Charmm22-CMAP/Tip3p)− to computing the folding energetics of a set of two-state and three-state proteins that do or do not carry a bound cofactor. The proteins successfully computed are representative of the main protein structural classes, their sequences range from 84 to 169 residues, and their isoelectric points from 4.0 to 8.9. The devised approach enables accurate calculation of two essential magnitudes governing the stability of proteins −the changes in enthalpy and in heat capacity associated to protein unfolding−, which are used to obtain accurate values of the change in Gibbs free-energy, also known as the protein conformational stability. The method proves to be also suitable to obtain changes in stability due to changes in solution pH, or stability differences between a wild-type protein and a variant. The approach addresses the calculation by difference, a shortcut that avoids having to simulate the protein folding time, which is very often unfeasible computationally.

## Introduction

Proteins are very versatile biological molecules^1^ and thermodynamics can greatly help to understand how they fold and perform useful tasks^2,3^. Molecular Dynamics (MD) simulation has become a powerful tool to study protein folding and other related processes^4–12^. However, despite great efforts in developing algorithms and methods to enable longer and better sampled simulations and in improving the accuracy of force fields and water models, significant challenges remain^13^. On one hand, simulating the protein folding time (from microseconds up to tens of seconds or beyond) in explicit solvent remains inaccessible, except for small fast-folding proteins^5–7,9^. On the other, work on improving the accuracy of MD force fields seems to have focused on reproducing structural, dynamic and mechanistic aspects of protein behavior^14–18^, and has paid less attention to try to reproduce protein potential energy. One reason for this is the difficulty in obtaining accurate structural models of unfolded ensembles, which has prevented comprehensive studies of this side of the problem, making fine-tuning of the force field parameters difficult. The experimental limitations inherent in quantifying individual atomic interactions, and the massive cancellation of interactions that takes place in a protein folding reaction^19^ adds to the complexity of the goal^2^. All of the above has perhaps frustrated the interest of scientists in the use of MD simulations to quantitatively study protein thermodynamics, hindering progress in many applied fields, such as protein design^20^, drug design^21^, genetic interpretation^22^, protein engineering^23^ or cell engineering^24^.

Recently, we addressed this issue by carrying out accurate, quantitative calculations of conformational stability on two two-state model proteins (barnase and nuclease) through an all-atom MD-based approach^25^. The approach circumvents the simulation of the whole folding/unfolding time and it is based on separately simulating the two main conformations: the folded and unfolded states. The folded state is modeled starting from an experimentally determined structure that is conveniently solvated and sampled conformationally. The unfolded state is modeled and sampled from an ensemble of completely unfolded conformations generated by the ProtSA server^26^ that are similarly solvated. From the simulations, the enthalpy change of unfolding (ΔH_unf_) is calculated by difference (the unfolded minus the folded state enthalpy averages), while the heat capacity change at constant pressure (ΔCp_unf_) is obtained from the temperature dependence of the calculated enthalpy change. As a final step, the calculated thermodynamic quantities (ΔH_unf_ and ΔCp_unf_) are combined with the experimentally determined mid denaturation temperature (T_m_) to calculate the conformational stability of the protein (ΔG_unf_) as a function of temperature by means of the Gibbs-Helmholtz equation^27^. One goal of the approach was testing the ability of classical force fields, e.g. Charmm22-CMAP^18^ and AmberSB99-ILDN^17^ (or the more recently released Amber force field, AmberSB99-disp^16^, tuned to be used both with folded and disordered proteins) to yield accurate folding energetics by difference using systems solvated with explicit water. Therefore, the indicated force fields were combined with seven explicit water models, Tip3p^28^, Tip4p^28^, Tip4p-d^29^, Tip4-d-mod^16^, Tip5p^30^, Spc^31^, and Spc/E^32^.

Results obtained by setting short MD simulations (2-ns productive trajectories per replica) and the combinations of either Charmm22-CMAP or AmberSB99-ILDN with Tip3p allowed, for the two proteins indicated, to finely capture the energy balance between the numerous interactions established between protein and solvent atoms in both the native state and the unfolded ensemble^33,34^.

In this work, we generalize the described methodology using the most accurate combination of force field and water model found^25^ and a larger conformational sampling, and show the good accuracy obtained on a set of two-state, three-state, apo, holo, wild-type (WT) or mutated proteins relative to their corresponding experimentally determined thermodynamics. In addition to barnase^35,36^ and nuclease^37,38^ energetics (here calculated anew with higher precision^25^), we present the calculation for additional two-state proteins: Barley chymotrypsin inhibitor 2 (CI2, truncated variant^39,40^) and Phage T4 lysozyme^41^ (the WT and pseudo-WT variants), a three-state protein (apoflavodoxin from *Anabaena* PCC 7119^42,43–45^) for which the energetics involved in the two unfolding transitions (ΔG_unf(F-to-I)_, and ΔG_unf(I-to-U)_) are obtained, and a holoprotein (flavodoxin from *Anabaena* PCC 7119) containing the FMN cofactor. Furthermore, we evaluate the capability and limits of the approach to capture small stability changes or small differences between similar systems, e.g. those associated to mutation (ΔΔH_mut-nat_ and ΔΔG_mut-nat_), changes in pH (ΔΔH_pH1-pH2_ and ΔΔG_pH1-pH2_) or individual steps within a multi-state unfolding (ΔH_unf(F-to-I)_, ΔCp_unf(F-to-I)_, ΔG_unf(F-to-I)_ or ΔH_unf(I-to-U)_, ΔCpunf(I-to-U) and ΔG_unf(I-to-U)_). The method requires a reliable structural model for the folded conformation, which sometime is not available. Hopefully, advances in high resolution protein models^46,47^ will allow the application of the protocol to the entire proteome.

### Target Proteins and Case Studies

#### Barnase from *B. amyloliquefaciens* and nuclease from *S. aureus*

The 110- and 149-residue proteins barnase^14,48–57^ and nuclease^58–61^ (C-terminal fragment), respectively, are well characterized two-state proteins, as summarized in Galano-Frutos et.al.^25^, where their folding energetics were first calculated. Here, the two-state unfolding energetics of WT barnase and nuclease are again determined using the present computational approach and the reported effect of pH on nuclease stability is also analyzed (see **Table 1**).

**Table 1.**
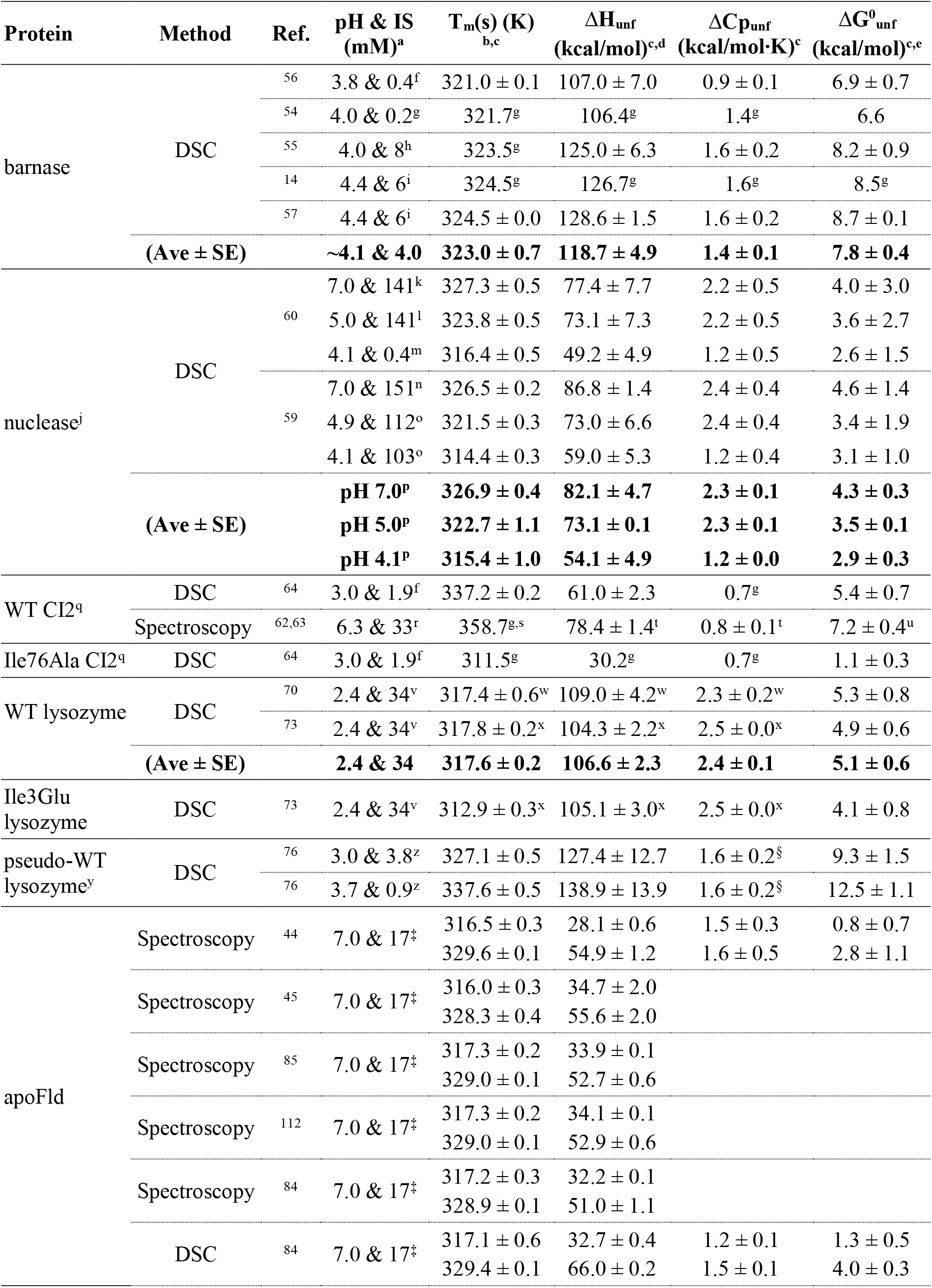
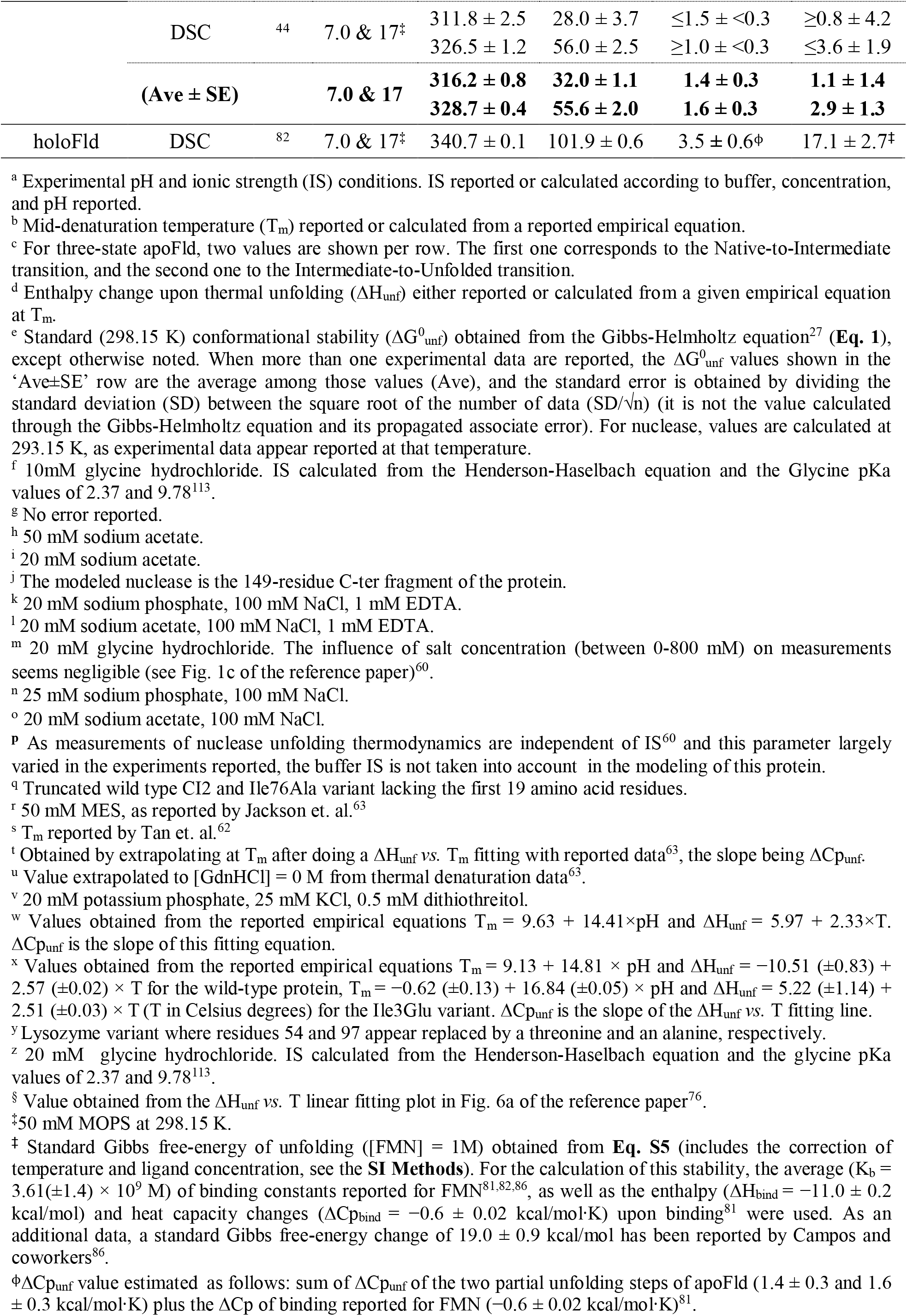
Experimental thermal unfolding data.

#### CI2 from barley seeds

CI2^39,40^ is a small, 84-residue, globular serine proteinase inhibitory protein extensively studied and reported to fold in a two-state manner under thermal conditions^62–64^. Its 19-residue N-terminal tail is completely unstructured^65,66^. We focus here on a truncated form of CI2 lacking the unstructured N-terminal tail because the structure of the full length protein is not available and because it has been shown that the tail does not contribute to the protein stability^63,64^. Thermodynamic data for the WT CI2 truncated form and for a broad set of point mutants analyzed under different solvent conditions (varying in pH and ionic strength) are available^62–64^ (**Table 1**). The CI2 variant Ile76Ala which, relative to WT in identical solvent, shows a significantly lower unfolding enthalpy change and a large destabilization^64^ (**Table 1**) has been selected to evaluate the feasibility of the approach to calculate the effect of single amino acid replacements on protein stability. On the other hand, the significantly different thermodynamics quantities reported for WT CI2^64^ under pH 3.0 and 6.3 (see **Table 1**), invited us to also evaluate the sensibility of the method to solvent effects.

#### Phage T4 lysozyme

T4 endolysin (lysozyme)^41^ is a two-domain, 164-residue globular protein that has also been subject of extensive study, and widely used to investigate the role of hydrophobic interactions in protein structural stabilization^67–69^. The thermal stability of the WT protein and many variants thereof have been measured^70–73^. Over 500 X-ray structures of T4 lysozyme –including those of an engineered pseudo-lysozyme (see below) and many variants thereof– have been obtained under a variety of experimental conditions (buffer, pH, ionic strength)^74^. WT lysozyme carries two cysteine residues at positions 54 and 97. To ease experimental work on the protein, a Cys54Thr/Cys97Ala variant (termed pseudo-WT lysozyme) has often been studied. WT and pseudo-WT lysozymes^75^ slightly differ in structure and thermodynamics^70–73,76^ (**Table 1**). For the sake of testing the method, the energetics of these two lysozyme variants are calculated. Besides, the energetics of the non-pseudo lysozyme variant, Ile3Glu^73^, is addressed as a further attempt to capture the effect of single amino acid replacements, and the pseudo-WT lysozyme^76^ is simulated in different solvent conditions (different pH values) to assess, as with nuclease and CI2, whether the method can capture pH-related effects on protein stability (**Table 1**).

#### *Anabaena* flavodoxin (Fld)

Fld^42,77^ is a 169-residue protein that carries electrons from photosystem I to ferredoxin-NADP^+^ reductase^78,79^ in the cyanobacteria *Anabaena* PCC 7119. Fld capability to transfer electrons is conferred by the presence of a molecule of non-covalently bound FMN cofactor. Reversible removal of the cofactor from the holoprotein (holoFld) leads to the apo form (apoFld). Fld has been widely studied to investigate protein/cofactor interactions^80,81^, as well as non-native protein conformations^44,82–85^. While apoFld thermal unfolding equilibrium is three-state^43–45^, binding of FMN greatly stabilizes the complex so that holoFld unfolds following a two-state mechanism^82,86^. A detailed picture of Fld folding and binding thermodynamics is available^43–45,81,82,84–87^. The reasonably high enthalpy and heat capacity changes (**Table 1**) of the two apoFld unfolding transitions, folded to intermediate (F-to-I) and intermediate to unfolded (I-to-U), together with the availability of a representative structure of its intermediate conformation^84^ made us select this protein to test the simulation approach for the calculation of unfolding energetics in three-state proteins.

## Results

### Energetics of two-state proteins: barnase, nuclease, CI2 and lysozyme unfolding

The thermal unfolding of barnase, nuclease, CI2 and lysozyme at equilibrium has been described to follow a two-state mechanism. Accordingly, we have calculated their unfolding energetics using the general workflow (see **Methods** and **Figure 1**), where the numbers of simulated replicas of the folded state and of simulated structures in the unfolded ensemble has been increased relative to the initial formulation of the method^25^.

**Figure 1.**
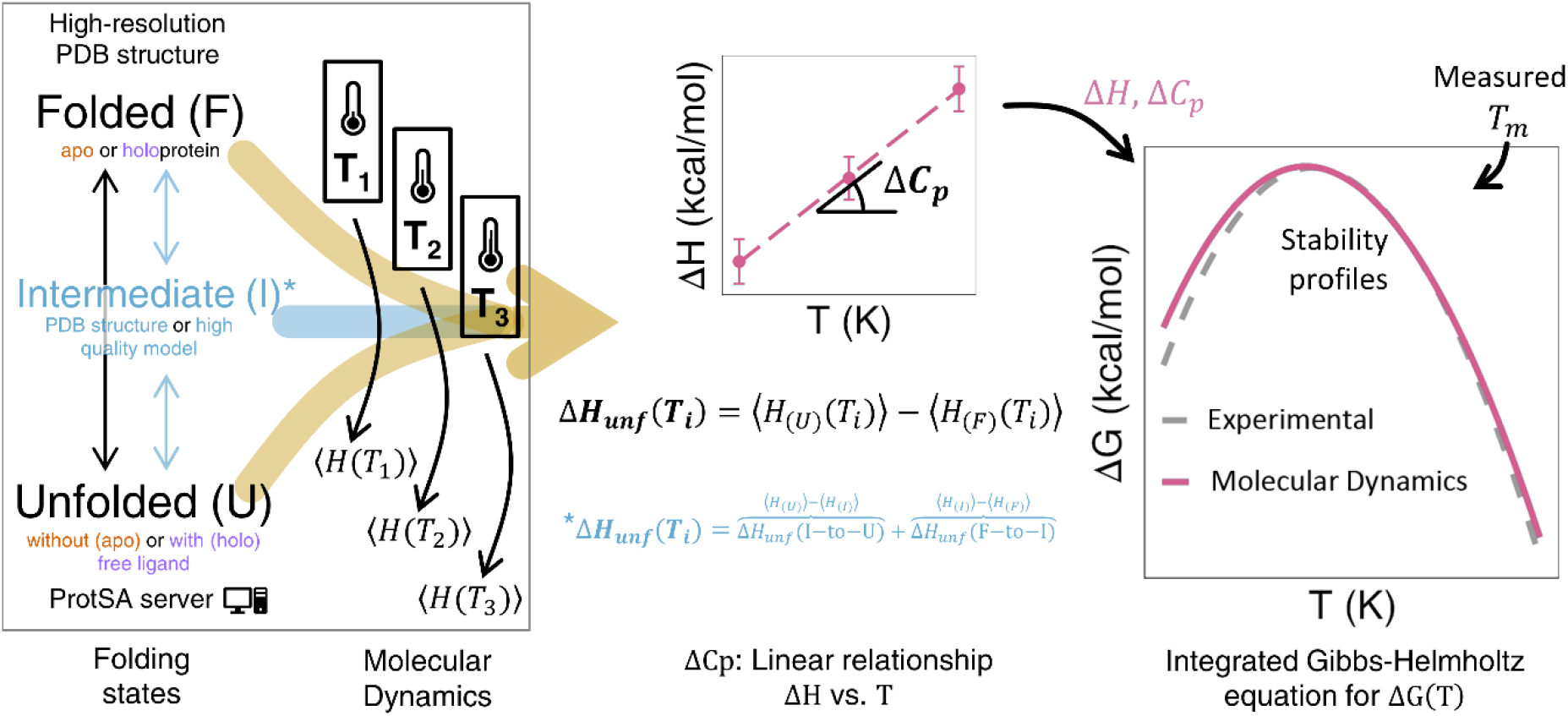
General workflow of the devised MD-based approach. The enthalpy of simulation boxes containing either folded (H_apo(F)_ or H_holo(F)_) or unfolded (H_apo(U)_ or H_apo(U)+cofactor_ in holoproteins) protein (or a structure representative of the intermediate state, H_apo(I)_) are directly computed and averaged from Molecular Dynamics simulations. The unfolding enthalpy change (ΔH_unf_) is obtained as the difference. The simulations are performed at three temperatures and the change in heat capacity (ΔCp_unf_) is obtained as the slope of a linear plot of enthalpy change versus temperature. These two magnitudes are combined with the experimental T_m_ of the protein to calculate the conformational stability using the Gibbs-Helmholtz equation. For holoproteins, a similar equation, **Eq. S5** in the **Supplementary Material**, is used that applies a correction to Gibbs free-energy to account for the ligand concentration, and uses the van’t Hoff approximation to describe the temperature dependence of the binding constant, K_b_(T). The number of water molecules and ions added to the folded, unfolded (or intermediate, if applicable) boxes must be identical. Forty (40) replicas of the folded box (normally built from a high-resolution PDB structure) and one hundred (100) of the unfolded box built from a filtered sample of completely unfolded conformations generated from ProtSA server^26^ are simulated. For intermediate states, 100 simulation replicas can be built from a representative structural ensemble. For holoproteins, the unfolded box is built by placing an unfolded protein molecule generated with ProtSA and one molecule of cofactor at a given minimum distance of the protein. The rest of general details can be found in **Methods** and in panels **a** of **Figures 2-4** and **Extended Data Figures 1-4**.

Barnase is simulated (**Figure 2a**) at pH ∼4.1 (see **Table 1** and **SI Table 2**) in solvating conditions similar to those used in the experimental measurements reported and also to those previously used to model the protein^25^. In the previous modelling, a reasonable agreement was found between experimental and calculated data. Here, the calculated values of ΔH_unf_(T_m_), ΔCp_unf_ and ΔG_unf_ at 298.15 K (ΔG^0^ _unf_) obtained with the extended sampling (110.4 ± 3.1 kcal/mol, 1.0 ± 0.1 kcal/mol·K and 7.5 ± 1.2 kcal/mol, respectively, **Table 2**) agree very well with the averaged experimental energetics (118.7 ± 4.9 kcal/mol, 1.4 ± 0.1 kcal/mol·K and 7.8 ± 0.4 kcal/mol, **Table 1**). Due to this fine agreement, the experimental and calculated temperature dependencies of ΔG_unf_ (stability curves), excess heat capacity (thermograms), and state fractions (**Figure 2b-d)** nearly coincide. The agreement between calculated and experimental magnitudes is better than that obtained in the previous calculation^25^ (92.3 ± 5.7 kcal/mol, 0.9 ± 0.1 kcal/mol·K and 6.5 ± 0.8 kcal/mol, respectively) that was performed using the same conditions and workflow but with a smaller sampling.

**Figure 2.**
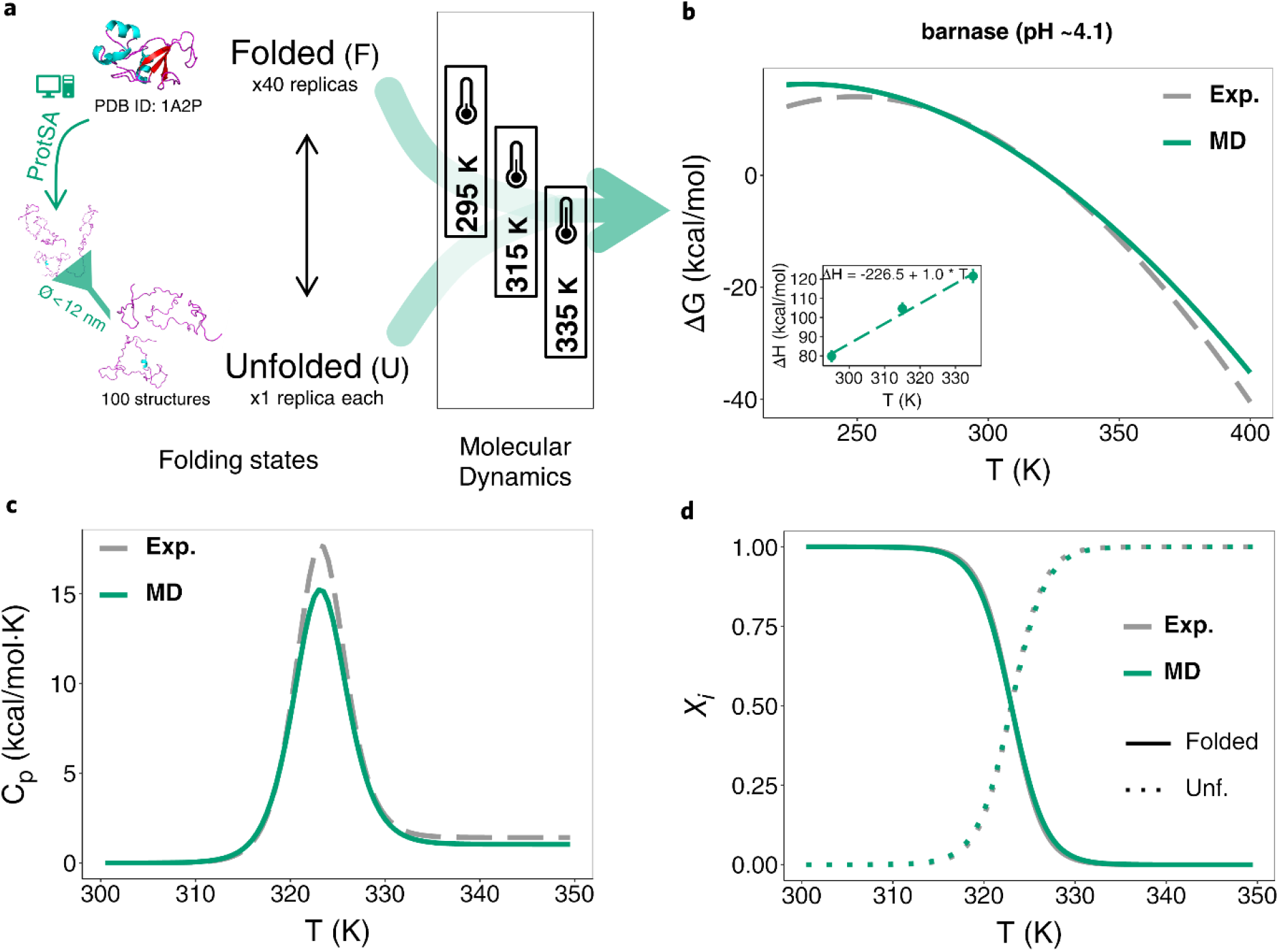
Simplified MD-based scheme and comparison with experimental results for a two-state protein example: barnase. **a)** The protein models, the number of structures (unfolded) and replicas (folded) simulated, the diameter cutoff used to filter too-elongated unfolded structures obtained from ProtSA^26^ (left), and temperatures selected for the MD-based calculation (Charmm22-CMAP) of thermodynamics of barnase. **b-d)** Temperature profiles of ΔG_unf_, Cp_unf_ and protein molar fractions (*χ*_*i*_) (*in silico* vs. experimental), respectively, obtained for barnase simulated at pH ∼4.1. Inset in **b** depicts the calculated ΔH_unf_ vs. T linear plot with the fitted equation (the slope being ΔCp_unf_) obtained from the MD simulations. The color-coding is indicated in the legends of the panels.

**Table 2.**
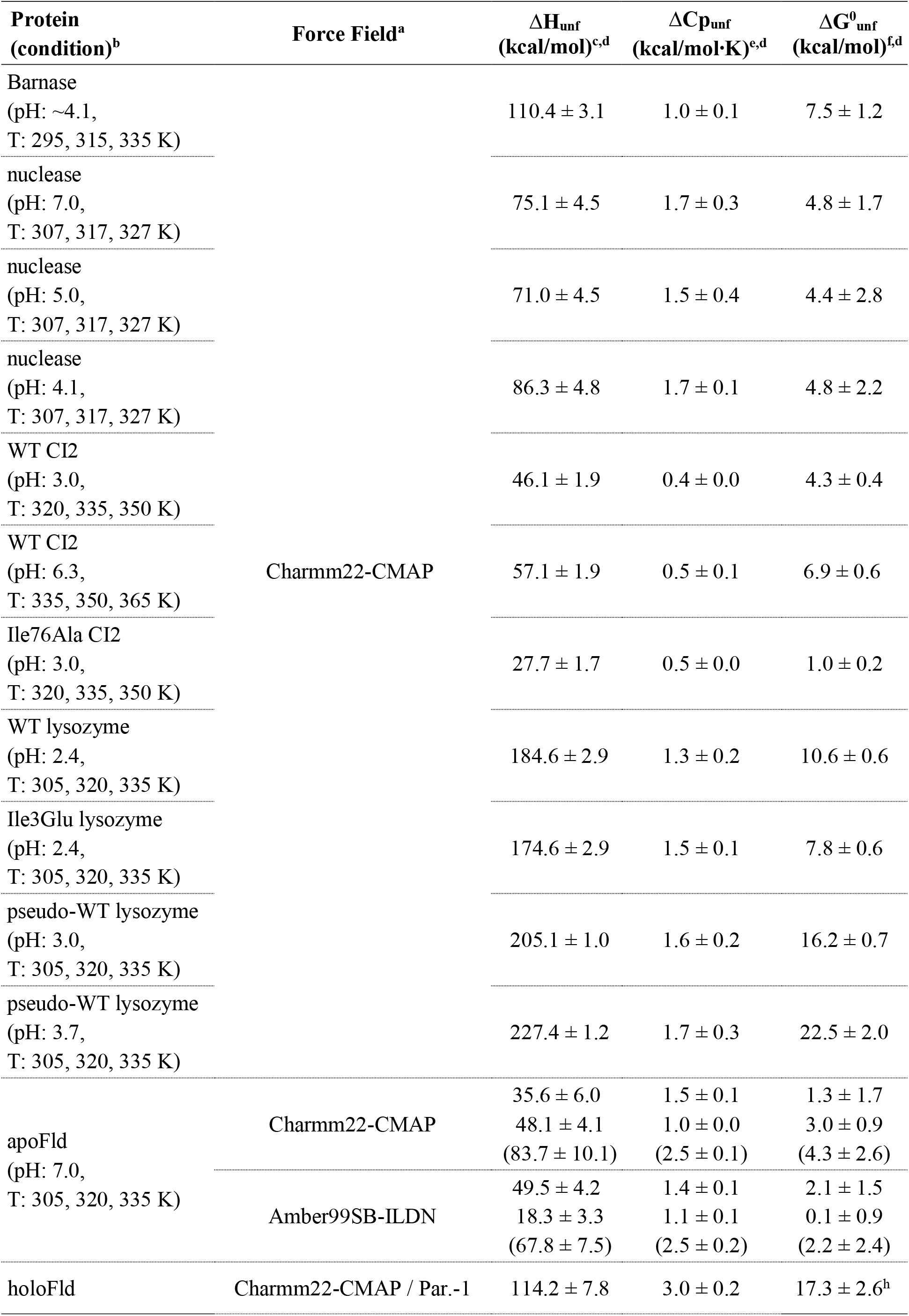
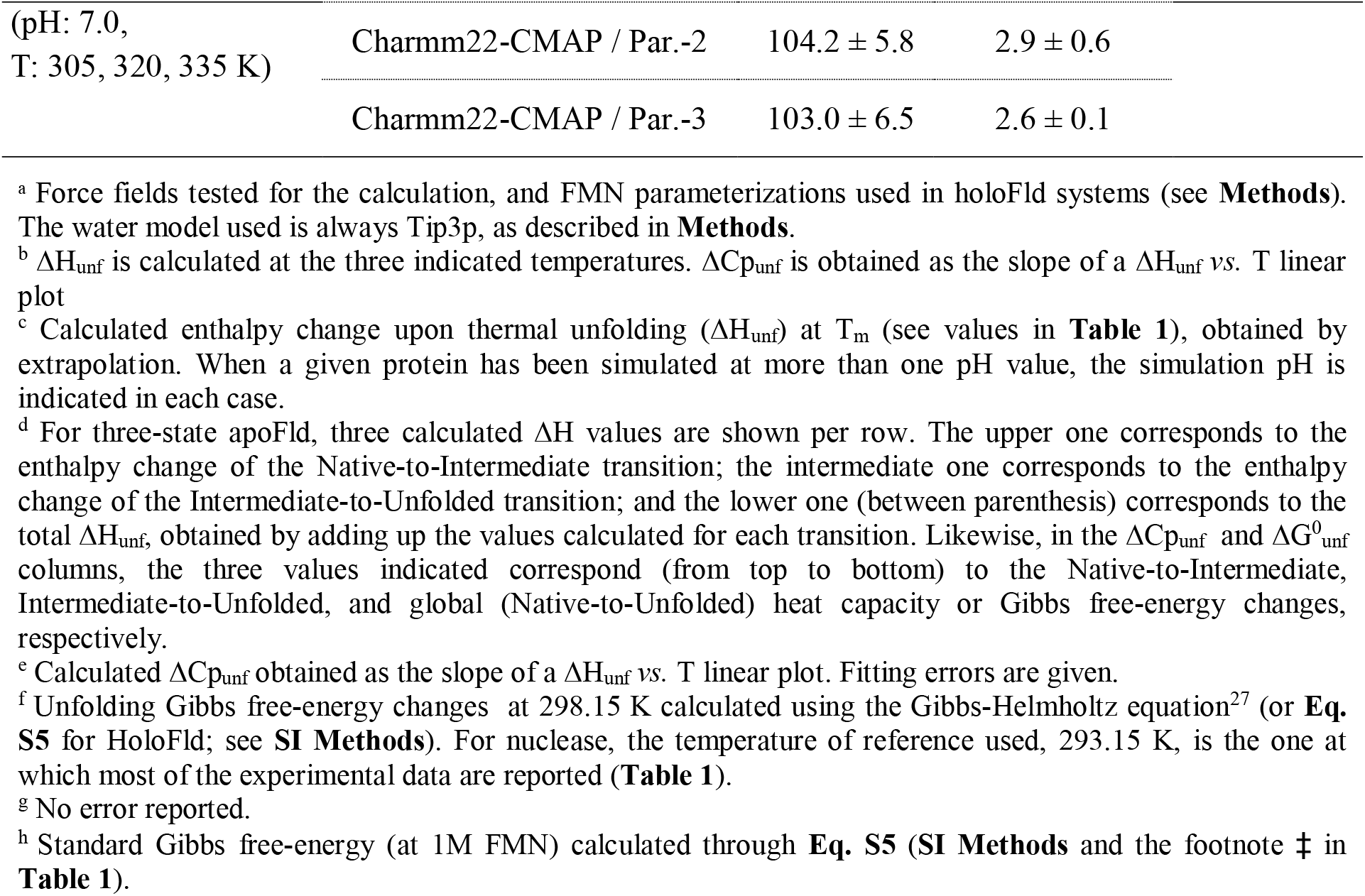
Calculated thermal unfolding energetics from MD simulations.

Nuclease unfolding thermodynamic data are available over a range of pH (from 3 to 8.5) and solvating conditions^59,60^. WT nuclease is simulated here (**Extended Data Figure 1a**) at three pH values: 7.0, 5.0 and 4.1 (see solvating conditions and protonation states in **Table 1** and **SI Table 2**). At pH 7.0, the calculated ΔH_unf_(T_m_) (75.1 ± 4.5 kcal/mol), ΔCp_unf_ (1.7 ± 0.3 kcal/mol·K) and ΔG_unf_ (4.8 ± 1.7 kcal/mol) at 293.15 K (the temperature at which experimental data are mostly reported for this protein, **Table 2**) match very well the averaged experimental values (82.1 ± 4.7 kcal/mol, 2.3 ± 0.3 kcal/mol·K and 4.3 ± 0.3 kcal/mol, respectively, **Table 1**). This excellent agreement is reflected, as seen above for barnase, in the fine correspondence between the experimental and calculated Gibbs free-energy difference, excess heat capacity, and molar fractions temperature dependencies (**Extended Data Figure 1b-d**). The second solvating condition simulated for nuclease reproduces a protonation scheme previously used^25^, corresponding to pH 5.0. Under this condition, our simulations yield calculated energetics (ΔH_unf_(T_m_) = 71.0 ± 4.5 kcal/mol, ΔCp_unf_ = 1.5 ± 0.4 kcal/mol·K and ΔG ^293.15K^_unf_ = 4.4 ± 2.8 kcal/mol, **Table 2**) that matches fairly well the mean experimental values (73.1 ± 0.1 kcal/mol, 2.3 ± 0.1 kcal/mol·K and 3.5 ± 0.1 kcal/mol, respectively, **Table 1** and **Extended Data Figure 1e-f**). The application here of a more exhaustive sampling reproduces similarly well the experimental results of nuclease compared to the implementation carried out with a smaller sampling^25^: calculated ΔH_unf_(T_m_) = 76.0 ± 8.1 kcal/mol, ΔCp_unf_ = 1.8 ± 0.1 kcal/mol·K and ΔG^293.15K^_unf_ = 4.6 ± 1.4 kcal/mol, respectively^25^. Thus, nuclease stability is accurately calculated in the 5.0-7.0 pH range. However, at lower pH, pH 4.1, the method overestimates ΔH_unf_(T_m_) and ΔCp_unf_, which leads to less accurate calculated stability (4.8 ± 2.2 kcal/mol, **Table 2** versus a mean experimental value of 2.9 ± 0.3 kcal/mol, **Table 1** and **Extended Data Figure 1g-h**).

WT CI2 is simulated (**Extended Data Figure 2a**) at two pH values for which reliable experimental data are available (**Table 1** and **SI Table 2**). At pH 3.0, the calculated ΔH_unf_(T_m_) and ΔCp_unf_ values (46.1 ± 1.9 kcal/mol and 0.4 ± 0.03 kcal/mol·K, respectively) are a bit lower than the corresponding experimental ones (61.0 ± 2.3 kcal/mol and 0.72 kcal/mol·K). Notwithstanding, the calculated ΔG^0^_unf_ at this pH (4.3 ± 0.4 kcal/mol) virtually agrees within error with the experimental stability reported at 298.15 K (5.4 ± 0.7 kcal/mol). At pH 6.3, CI2 is more stable. At this pH, the experimental ΔH_unf_(T_m_) and ΔCp_unf_ values (78.4 ± 0.7 kcal/mol and 0.8 ± 0.1 kcal/mol·K, respectively) lead to a higher conformational stability (ΔG^0^_unf_ = 7.2 ± 0.4 kcal/mol). The increases in ΔH_unf_(T_m_) and ΔCp_unf_ values at pH 6.3 relative to pH 3.0 are captured by our simulations (calculated values at pH 6.3: ΔH_unf_(T_m_) = 57.1 ± 0.5 kcal/mol and ΔCp_unf_ = 0.5 ± 0.07 kcal/mol·K, respectively), as well as the increase in conformational stability (calculated ΔG^0^_unf_ = 6.9 ± 0.6 kcal/mol, in agreement with the experimental value of 7.2 ± 0.4 kcal/mol). To assess the capability of the simulation approach to detect changes in stability associated to point mutations, we compute the energetics of the CI2 variant Ile76Ala at pH 3.0 and compared it to that of WT CI2 at the same pH. Substitution of the bulky WT isoleucine residue by alanine creates a cavity that severely destabilizes the folded structure of the mutant. The reduced stability of Ile76Ala CI2 compared to WT is evidenced in its experimental unfolding energetics (ΔH_unf_(T_m_) = 30.2 kcal/mol, ΔCp_unf_ = 0.7 kcal/mol·K and ΔG^0^_unf_ = 1.1 ± 0.3 kcal/mol, **Table 1**), which is accurately obtained from our simulations: calculated ΔH_unf_(T_m_) = 27.7 ± 1.7 kcal/mol, ΔCp_unf_ = 0.5 ± 0.01 kcal/mol·K and ΔG^0^_unf_ = 1.0 ± 0.2 kcal/mol (**Table 2**). Thus, the simulation workflow allows to capture the experimental observations that: 1) WT CI2 is stabilized by raising the pH from 3.0 to 6.3 (experimental ΔΔG_unf(pH3→pH6.3)_ = +1.8 ± 1.1 kcal/mol; calculated value = +2.5 ± 1.2 kcal/mol) and 2) WT CI2 is severely destabilized by replacing Ile76 by Ala (experimental ΔΔG^0^_unf(WT→I76A)_ = −4.3 ± 1.0 kcal/mol; calculated value = −3.3 ± 0.6 kcal/mol). Experimental and calculated stability curves, thermograms, and state fractions of WT (pH 3.0), WT (pH 6.3) and Ile76Ala CI2 mutant (pH 3.0) are compared in **Extended Data Figure 2b-h**. A good agreement between calculated and experimental data can be observed that is particularly remarkable for the Ile76Ala CI2 variant (**Extended Data Figure 2g-h**).

Lysozyme is simulated (**Extended Data Figure 3a**) at pH 2.4 (WT and Ile3Glu mutant) and at pH 3.0 and 3.7 (pseudo-WT, **Extended Data Figure 4a**). The experimental ΔCp_unf_ is calculated accurately for the pseudo-WT but underestimated for the WT. For the four simulated lysozyme variants or pH condition (**Table 1**), the calculated ΔH_unf_ values (**Table 2**) clearly overestimate the corresponding experimental ones (**Table 1**). As a consequence, the stabilities calculated at 298.15 K also overestimate the experimental values, and the stability versus temperature dependencies (**Extended Data Figures 3b-f** and **4b-f**) do not match the calculated ones. Thus, the actual lysozyme stabilities are not correctly calculated by this MD-based approach (possible reasons are indicated in the **Discussion** section). Still, both the lower stability of the Ile3Glu mutant relative to WT at pH 2.4 (ΔΔG^0 unf(WT→Ile3Glu)^ = −1.0 kcal/mol), and the higher stability of pseudo-WT at pH 3.7 compared to pH 3.0 (ΔΔG_unf(pH3.0→pH3.7)_ = +3.3 kcal/mol) are qualitatively captured (−2.8 and +6.5 kcal/mol, respectively).

### Energetics of a three-state protein: apoFld unfolding

ApoFld thermal unfolding is three-state, with a well-defined intermediate accumulating at equilibrium with the folded and unfolded conformations. The unfolding enthalpy changes of the sequential partial unfolding equilibria (F-to-I and I-to-U) are separately calculated using the general workflow (**Figure 1**) from the structures or ensembles representing the states involved in each transition (**Figure 3a**). The calculated enthalpy changes of the two unfolding transitions, ΔH_unf(F-to-I)_ = 35.6 ± 6.0 and ΔH_unf(I-to-U)_ = 48.1 ± 4.1 kcal/mol (**Table 2**), are in excellent agreement with the corresponding experimental enthalpies of 32.0 ± 1.1 and 55.6 ± 2.0 kcal/mol (**Table 1**). The heat capacity changes calculated for each partial unfolding step, ΔCp_unf(F-to-I)_ = 1.5 ± 0.1 and ΔCp_unf(I-to-U)_ = 1.0 ± 0.0 kcal/mol·K, respectively (2.5 ± 0.1 kcal/mol·K for the global transition, **Table 2**) are also in fair agreement with the experimental values of 1.35 ± 0.3 and 1.55 ± 0.3 kcal/K·mol, respectively (2.9 ± 0.6 kcal/mol·K for the global transition, **Table 1**). From these calculated data and the experimental T_m_s of the two transitions (**Table 1**), the Gibbs free-energy changes of the individual and global unfolding transitions are calculated at 298.15 K, with the Gibbs-Helmholtz equation^27^. A fine correspondence between the calculated values, ΔG^0^_unf(F-to-I)_ = 1.3 ± 1.7 kcal/mol, ΔG^0^ _unf(I-to-U)_ = 3.0 ± 0.9 kcal/mol and ΔG^0^ _unf(F-to-U)_ = 4.3 ± 2.6 (**Table 2**), and the corresponding experimental ones, 1.1 ± 1.4, 2.9 ± 1.3 and 4.0 ± 2.7 kcal/mol, is observed. This outstanding correspondence between the calculated and the experimentally determined apoFld thermal unfolding thermodynamics can be also visually assessed from the stability curves^27^, thermograms, and folded/intermediate/unfolded state-fractions (**Figure 3b-d**).

**Figure 3.**
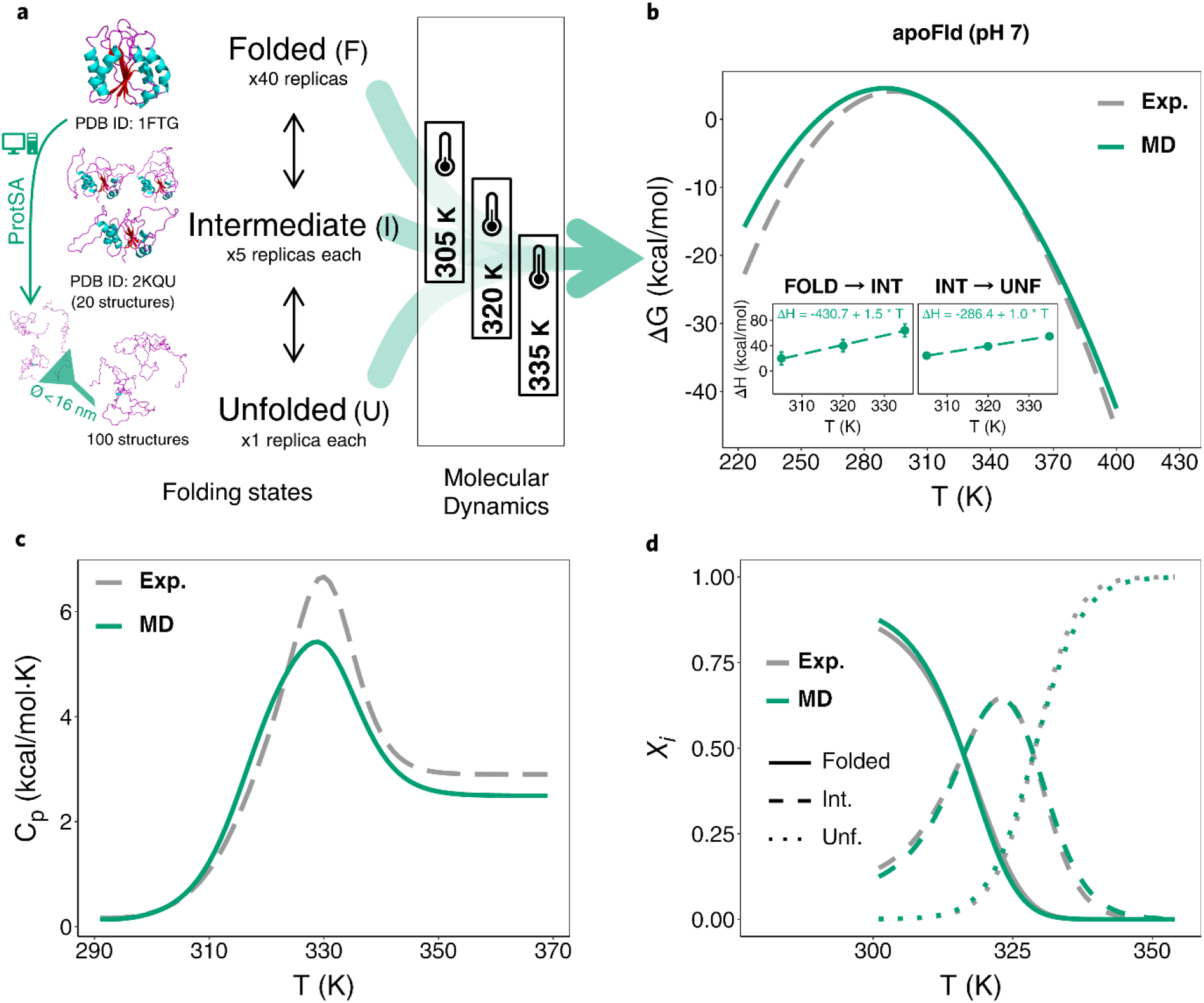
Simplified MD-based scheme and comparison with experimental results for a three-state protein example: apoFld. **a)** Protein models, number of structures (unfolded) and replicas (folded) simulated, diameter cutoff used to filter too-elongated unfolded structures obtained from ProtSA^26^ (left), and temperatures selected for the MD-based calculation (Charmm22-CMAP) of apoFld thermodynamics. **b-d)** Temperature profiles of the global stability (ΔG_unf_ = ΔG_unf(F-to-I)_ + ΔG_unf(I-to-U)_), Cp_unf_ and *χ*_*i*_, respectively (*in silico* vs. experimental). Inset in **b** depicts linear plots of calculated ΔH_unf_ from the MD simulations *vs*. T, with the fitted equation (the slope being ΔCp_unf_) obtained. The color-coding is indicated in the legends of the panels.

For the sake of comparison, an otherwise identical calculation of apoFld thermal unfolding thermodynamics has been carried out using the Amber99SB-ILDN force field instead of Charmm22-CMAP. Although accurate heat capacity changes (ΔCp_unf_) are calculated for the two equilibria with Amber99SB-ILDN (1.4 ± 0.1 and 1.1 ± 0.1 kcal/K·mol, respectively, **Table 2**), the calculated enthalpy changes (**Table 2**) do not agree well with the experimental values (**Table 1**), which results in less accurate calculations of the sequential Gibbs free-energy changes (**Table 2**), compared to those obtained with Charmm22-CMAP. A better agreement of Charmm22-CMAP thermodynamics calculations with experimental values was already reported for barnase and nuclease^25^.

### Energetics of a holoprotein: holoFld unfolding

The calculation of the thermal unfolding energetics of a holoprotein (a protein carrying a non-covalently bound cofactor) is performed as described in **Methods** and depicted in **Figure 4a**. The results obtained for the three different FMN parameterizations tested are presented in **Table 2** and **Figure 4b**. The ΔH_unf_ calculated for holoFld with any of the three setups (ranging from 103.0 ± 6.5 to 114.2 ± 7.8 kcal/mol) are in fair agreement with the experimental value reported by Lamazares and coworkers^82^ from DSC measurements (101.9 ± 0.6 kcal/mol, **Table 1**).

**Figure 4.**
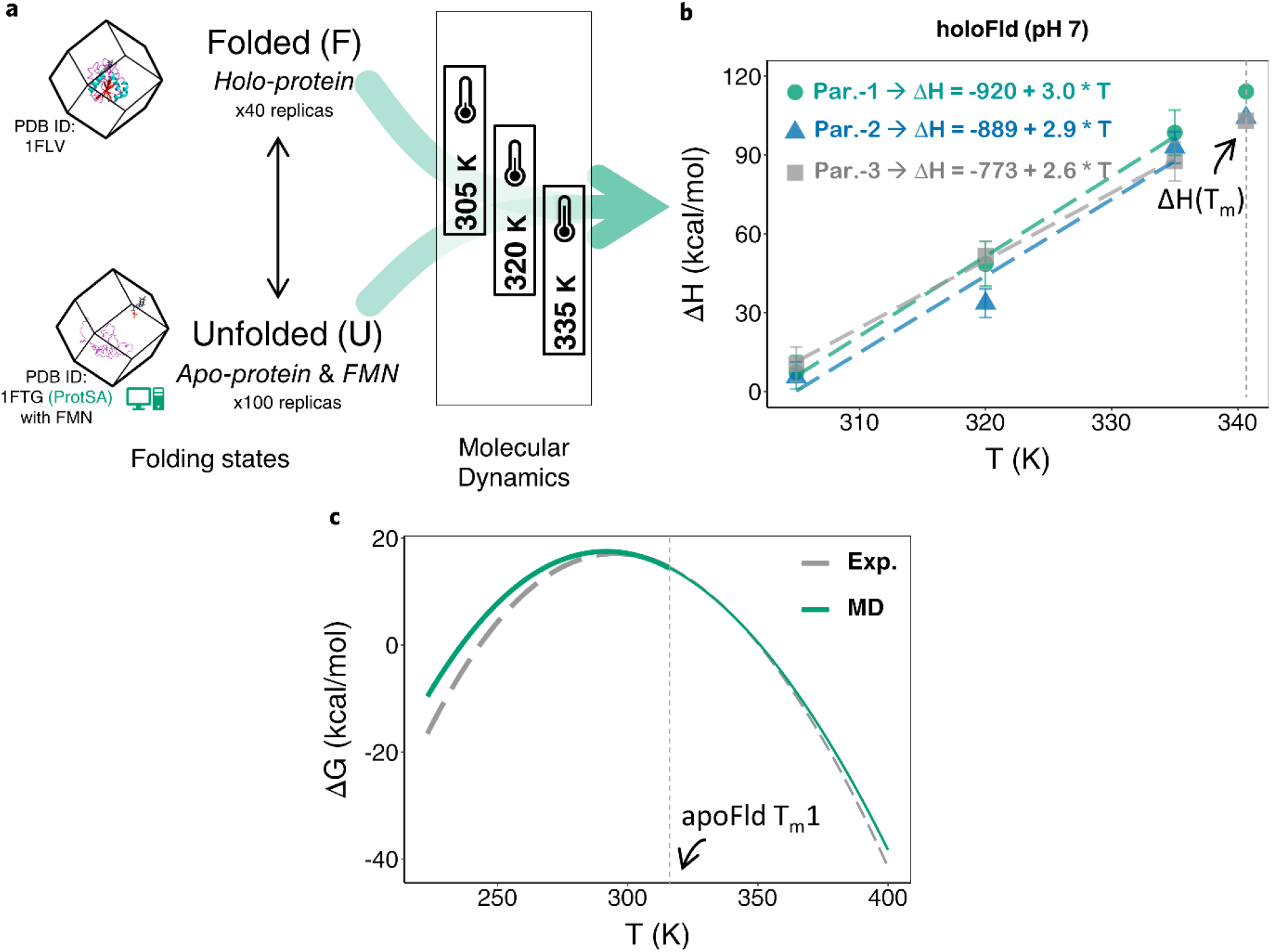
Simplified MD-based scheme and comparison with experimental results for a holoprotein example: holoFld. **a)** Protein and cofactor models placed in the simulation boxes, folding states, number of structures (unfolded) and replicas (folded) simulated, and temperatures selected for the MD-based calculation (Charmm22-CMAP) of holoFld thermodynamics. **b)** Calculated ΔH_unf_ vs. T linear plots, with the fitted equations (slopes are the respective ΔCp_unf_) obtained for the three FMN parameterizations tested. Extrapolated ΔH_unf_ values at T_m_ (340.7 K) are indicated over the vertical dashed line placed at this temperature. **c)** Temperature profiles of ΔG_unf_ (*in silico* vs. experimental) obtained from **Eq. S5**. Curves appear depicted with finer lines beyond the first T_m_ of the apoprotein (316.2 K, **Table 1**, vertical dashed line) to indicate –as described in **Results**– that in this region the ΔG_unf_ values calculated are not reliable. The fact that at 298.15 K the calculated stability of HoloFld (17.3 ± 2.6 kcal/mol) agrees within error with the stability measured from thermal unfolding curves (19.0 ± 0.9 kcal/mol)^86^ seems to validate the accuracy of the profiles in the range of temperatures below the apoprotein T_m_1.

The holoFld ΔCp_unf_ has not been reported, but an estimation can be done by adding the reported value for FMN dissociation (ΔCp_diss_ = −ΔCp_bind_ = 0.6 ± 0.0 kcal/mol·K) to the apoFld ΔCp_unf_ (2.9 ± 0.6 kcal/mol·K, **Table 1**). Thus, holoFld ΔCp_unf_ is estimated to be 3.5 ± 0.6 kcal/mol·K. Our calculated holoFld ΔCp_unf_ values are reported in **Table 2** for the three FMN parameterizations above referred and depicted as the slope of fitting lines in **Figure 4b**. The values obtained with either FMN Par.-1 or FMN Par.-2 (3.0 ± 0.2 and 2.9 ± 0.6 kcal/mol·K, respectively) agree within experimental error, and that obtained with FMN Par.-3 (2.6 ± 0.1 kcal/mol·K) while lower, is still above the value previously calculated for apoFld (2.5 ± 0.1 kcal/mol·K, **Table 2**), in agreement with the observed positive value of ΔCp for FMN dissociation.

The stability of holoFld at 298.15 K is obtained through **SI Eq. 5** (see the derivation in **SI Methods**). To the apoprotein Gibbs free-energy, **SI Eq. 5** applies a correction due to the ligand concentration, incorporating the van’t Hoff approximation^88^ to account for the temperature dependence of the binding constant, K_b_(T). Thus, **Eq. S5** is not based on the thermodynamics derived from the holoFld simulations (ΔH^holo^_unf_, ΔCp^holo^_unf_) but on those of the apoprotein (ΔH^apo^_unf_, ΔCp^apo^_ung_) plus the cofactor energetics. Using **SI Eq. 5**, the ΔG^298.15K^_unf_ value calculated (17.3 ± 2.6 kcal/mol, **Table 2**) is in close agreement with the experimental value (17.1 ± 2.7 kcal/mol, **Table 1**) similarly obtained with **SI Eq. 5** using experimental ΔH^apo^_unf_ and ΔCp^apo^ _unf_ data. Importantly, the calculated ΔG ^298.15K^ _unf_ also matches, within error, the experimental stability of holoFld directly obtained from thermal unfolding curves (19.0 ± 0.9 kcal/mol)^86^.

The van’t Hoff approximation to model the temperature dependence of binding constant^88^ −therefore the correction to stability due to the ligand concentration− should work fine as long as the conformation of the protein binding site does not change significantly. However, as this will not be the case at temperatures where the apoprotein begins to unfold, we consider the ΔG_unf_ temperature profile of the holoprotein (**Figure 4c**) not to be reliable beyond the first mid denaturation temperature (T_m_1) of the apoprotein (316.2 K in the case of apoFld).

## Discussion

### Overall accuracy of calculated energetics

The devised MD-based workflow allows the calculation of ΔH_unf_, ΔCp_unf_, and ΔG_unf_, the main thermodynamic magnitudes governing the stability of proteins. The overall accuracy of the method can be assessed from lineal plots of calculated versus experimentally determined values of each of those magnitudes.

The primary outcome is the unfolding enthalpy change of the proteins investigated. With the exception of lysozyme (simulated in four conditions) and nuclease (when simulated at low pH, pH 4.1), which are clear outliers, the linear plot (**Figure 5a**) can be fitted to a straight line with ordinate close to zero, slope close to unity (0.95) and a correlation of R^2^ = 0.93. The fitting includes the data from ten simulated systems (barnase, nuclease at two pH values, two partial unfolding equilibria as well as the whole transition of three-state apoFld, CI2 at two pH values plus one mutant, and holoFld) spanning a range of ΔH_unf_ values from 30 to 120 kcal/mol. It is thus clear that ΔH_unf_ can be accurately calculated using the approach.

**Figure 5.**
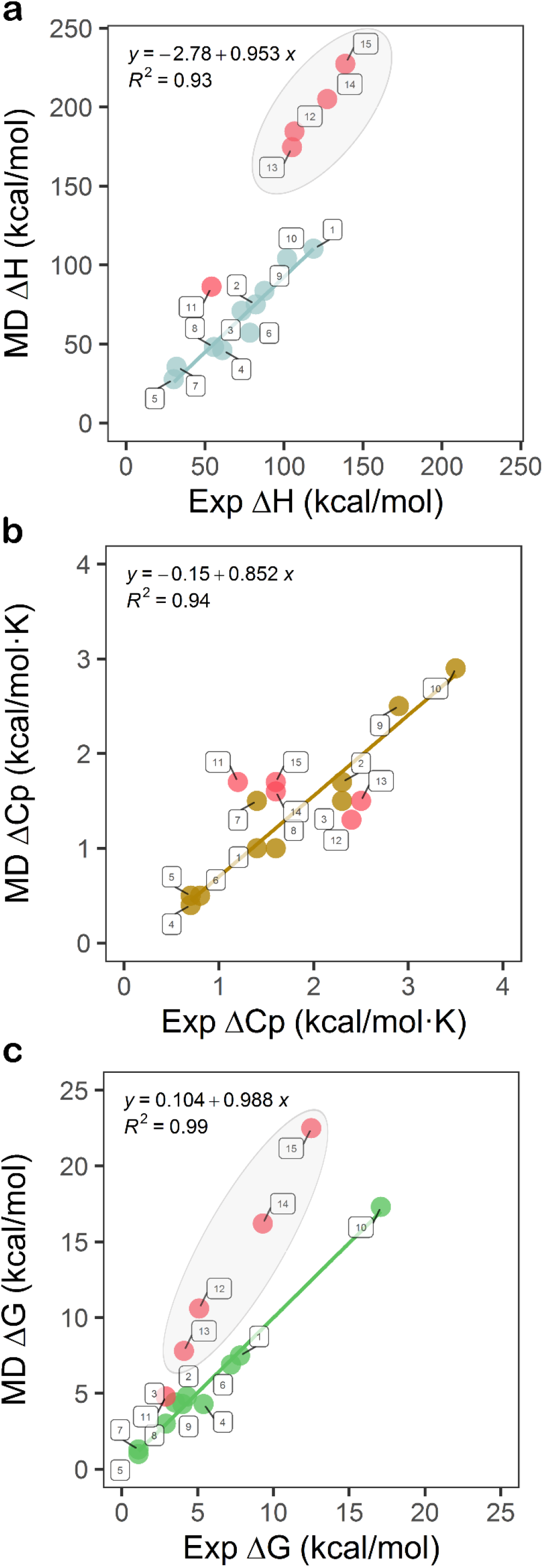
Global assessment of the approach for calculation of unfolding thermodynamics with Charmm22-CMAP/Tip3p. **a)** Scatter plot of MD-calculated vs. experimental ΔH_unf_ for the set of proteins simulated (including different solvating conditions and variants). The linear fit shown in this panel (also in panels **b** and **c**) was performed over the following ten systems: barnase at pH ∼4.1 (dot number 1 in legend), nuclease at pH 7.0 (2) and pH 5.0 (3), WT CI2 at pH 3.0 (4), Ile76Ala CI2 at pH 3.0 (5), WT CI2 at pH 6.3 (6), apoFld(F-to-I) (7), apoFld(I-to-U) (8), apoFld(F-to-U) (9) and holoFld(FMN Par.-2) (10). The fitting equation and the square Pearson correlation coefficient are given. **b)** Scatter plot and linear fit of MD-calculated vs. experimental ΔCp_unf_. **c)** Scatter plot and linear fitting of MD-calculated vs. experimental protein stability (ΔG_0_^unf^ at 298.15 K for all proteins except for nuclease that is compared at 293.15 K). Experimental values (*x*-axis) are the averages (or individual value in some cases) of data obtained from literature, as summarized in **Table 1**, while calculated values are those presented in **Table 2**. Red circles represent outliers (or cases treated as such, see the **Results and Discussion** section) not considered in the linear fitting, namely: nuclease at pH 4.1 (dot number 11 in legend), WT lysozyme at pH 2.4 (12), Ile3Glu lysozyme at pH 2.4 (13), pseudo-WT lysozyme at pH 3.0 (14) and pseudo-WT lysozyme at pH 3.7 (15). In panels **a** and **c**, the 4 outliers of lysozyme and pseudo-lysozyme systems are enclosed in a semi-transparent gray oval to visualize them as similar systems whose enthalpy change upon unfolding (ΔH_unf_) and protein stability (ΔG_unf_) are all overestimated by our simulations. Of the three setups tested for holoFld, the results obtained with FMN parameterization 2 (the most accurate one, see **Tables 1** and **2**) are depicted.

The second outcome is the unfolding heat capacity change (**Figure 5b**), which is also captured for those 10 protein systems well fitted in **Figure 5a**. The four lysozyme systems simulated (WT at two pH values, pseudo-WT and a variant of pseudo-WT), as well as nuclease at pH 4.1, fit worse than the other 10 systems. Albeit their calculated ΔCp_unf_ values do not differ too much from their experimental ones, they are also treated as outliers, for consistency. The linear fit with data from the 10 simulated systems also yields a straight line with ordinate close to zero, slope close to unity and a correlation of R^2^ = 0.94, indicating that the change in heat capacity of unfolding can be also calculated in an accurate manner using the approach. The range of ΔCp_unf_ values spanned goes from 0,6 to 3.5 kcal/mol·K.

The third outcome is the change in Gibbs free-energy upon unfolding (ΔG_unf_), i.e. the conformational stability of the protein. To derive it, our workflow combines the calculated enthalpy and heat capacity changes with experimental values of melting temperatures, using the Gibbs-Helmholtz equation (**Eq. 1**) for apoproteins, or an analogous equation for holoproteins (**SI Eq. 5**). As expected, in the linear plot of calculated versus experimentally determined stabilities **(Figure 5c)** lysozyme yields outliers as the high enthalpy changes calculated for this protein systems are carried over in the calculation of the stability. Although, nuclease at pH 4.1 is not a clear outlier in the stability representation, it is kept as such for consistency. The fitting of the calculated and experimental values for the other 10 systems simulated gives rise again to a straight line with close to zero intercept, close to unity slope and a high correlation of R^2^ = 0.99. It seems thus that conformational stability is also accurately calculated from first principles using the described simulation workflow. The range of Gibbs free-energies spanned in the plot goes from 1 to 17 kcal/mol.

### Applicability and limitations of the approach

The present MD-based workflow accurately calculates the protein changes in enthalpy, heat capacity and Gibbs free-energy upon unfolding and can also be used to compare the stability of a protein under different pH values, or to compare the stability of a wild-type protein with mutants thereof. According to our literature search, no similar approach for the calculation of protein folding energetics has been described, which precludes direct comparison with other methods. The systems successfully calculated here contain representatives of the main protein classes (mainly-alpha, mainly-beta, and alpha-beta)^89^, with sequences ranging from 84 to 169 residues, and isoelectric points from 4.0 to 8.9. They include proteins that undergo two- or three-state thermal unfolding, as well as proteins that do or do not carry a tightly bound cofactor. As a whole, these proteins offer a fair representation of natively folded proteins for which the unfolding process leads to fully unfolded conformations. As detailed thermodynamic studies on much larger proteins scarce, the approach has not been tested on large proteins. We foresee no reasons why the energetics of larger proteins cannot be calculated with similar accuracy using sufficient sampling, provided they exhibit fully unfolded conformations after heating. This is a requisite of our protocol, necessary to be able to build realistic models of the unfolded ensemble using ProtSA.^26^

However, for one of the proteins simulated, lysozyme, the calculations have consistently led to overestimated ΔH_unf_ values, which has translated in overestimated stability. In principle, the method could have failed due to insufficient quality of the models used to represent the lysozyme folded and unfolded conformations. This is unlikely as the folded structures have been solved in a highly experience lab^90^ and they get good marks when subjected to quality control with the MolProbity server^91^ (not shown). Besides, the model of the unfolded ensemble generated by ProtSA^26^ would be wrong if the lysozyme unfolded state were compact, but we have found no reports pointing to that. A different reason for the inaccurate lysozyme calculation may relate to small inaccuracies in force field parameters. Although the same force field has been used in lysozyme and in the successful calculations, it should be noticed that force field parameters are globally optimized and individual optimal performance of each of them cannot be taken for granted. In this respect, of all the systems simulated here, lysozyme stands out as the one containing the highest net (positive) charge (**Table S2**), only paralleled by the high net (positive) charge of nuclease in the single simulation condition (pH 4.1) where inaccurate results have also been obtained for this protein. It is possible that the discrepancy between calculated and experimental lysozyme unfolding magnitudes is related to insufficient tuning of coulombic treatment by the Charmm22-CMAP force field^18^, at least for lysine and arginine protonated side chains. In addition, some uncertainty in the protonation state of lysozyme carboxyl groups at the acidic pH of the simulations could also contribute to inaccuracy. Whatever the reason, the poorer performance of the method on lysozyme suggests it should be used with caution when highly positively charged proteins are simulated at acidic pH values. On the other hand, as proteins are rarely studied experimentally in basic pH conditions, we have not tested the performance of the method in alkaline solutions.

The sampling requirements of the protocol here presented are affordable with ordinary CPU- or GPU-clusters. The biggest, more challenging system here computed, 168-residue three-state apoFld (2.2 μs simulation: ∼3.0 ns per 40 folded, 100 intermediate and 100 unfolded replicas, each at three temperatures), takes 2-3 weeks of essentially unattended calculation on 32 cores (32 OpenMP threads/1 MPI task) of an Intel Xeon E5-2680v3-based cluster (170 CPUs arranged in 85 nodes, https://cesar.unizar.es/hpc/).

Although the described approach is based on a specific force field and water model, it suggests that current force fields are already close to capturing the complexity of protein folding energetics. We hope that our results will encourage further improvement of force fields and water models. Toward that goal, the described methodology constitutes an effective and efficient way to assess their ability to replicate the changes in energy that govern protein equilibria.

## Methods

### General MD-based workflow for calculation of unfolding energetics (ΔH_unf_, ΔCp_unf_ and ΔG_unf_) in apoproteins

The presented workflow (**Figure 1**) is based on that described in Galano-Frutos et.al.^25^. The current version relies on a higher sampling of the folded and unfolded states. Briefly, X-ray crystal structures with the highest resolution and sequence coverage are retrieved from the RCSB Protein Data Bank (https://www.rcsb.org/^92,93^, see PDB codes below) and taken as the starting structures for modeling the native (folded) state. When needed, the initial crystal structure is used to model the amino acid replacement leading to the mutant simulated (e.g. the CI2 Ile76Ala and lysozyme Ile3Glu variants).

Forty replicas of the folded structure are simulated, each consisting of a single protein molecule solvated with water molecules in a specified simulation box additionally containing, when required, ions (Na^+^ and/or Cl^−^). On the other hand, large unfolded ensembles (∼2000 structures) are generated from the protein sequence using the ProtSA server^26^ and a random sample of 100 unfolded structures is extracted to model the unfolded state (see **Figure 1** and panel **a** in **Figures 2-4** and **Extended Data Figures 1-4**). To avoid using too large simulation boxes, which may increase the simulation time as well as add noise to the results, the most extended conformations (∼10 %) are previously identified and removed as described^25^ (**SI Figure 1**). The selected 100 unfolded conformations are simulated in boxes containing one unfolded molecule and exactly the same number of water molecules, ions and cofactors –when it is the case− as in the corresponding boxes used to simulate the folded conformations of the same protein. For three-state proteins, in addition to the overall enthalpy change, those of the individual steps (F-to-I and I-to-U) can be obtained if the absolute enthalpy of an additional box containing one molecule of protein in the intermediate conformation and the same number of water and ion entities is calculated (see **Figure 1** and **Figure 3a**). To model the intermediate conformation, a suitable structural model is needed. In the three-state apoFld, a 20-model NMR ensemble previously described^84^ is used. In this case, five replicas are simulated for each of the 20 structures, totaling 100 replicas, the same number of unfolded conformations modeled (**Figure 3a**).

For each replica, a short 2-ns productive trajectory (see **SI Table 1**) is run and the individual time-averaged enthalpy (H^i^_F_, H^i^_U_ or H^i^_I_) retrieved. The individual enthalpies of replicas of the same conformational state (i.e. folded, unfolded or intermediate) are then ensemble-averaged to obtain the corresponding enthalpy to each folding state (⟨H_F_⟩, ⟨H_U_⟩ or ⟨H_I_⟩). Subsequently, the unfolding enthalpy change, ΔH_unf_, is calculated by difference, i.e. by subtracting the calculated ensemble-averaged enthalpy obtained from simulations of the folded state from the ensemble-averaged enthalpy obtained from simulations of the unfolded state: ΔH_unf_ = ⟨H_U_⟩ − ⟨H_F_⟩. For three-state proteins, enthalpy changes corresponding to the first unfolding transition (F-to-I) and the second one (I-to-U) are calculated likewise: ΔH_unf(F-to-I)_ = ⟨H_I_⟩ − ⟨H_F_⟩ and ΔH_unf(I-to-U)_ = ⟨H_U_⟩ − ⟨H_I_⟩ (**Figure 1**).

The calculation of the heat capacity change upon unfolding (ΔCp_unf_) relies on the linear dependency of ΔH_unf_ with temperature. For each protein, three not-distant temperatures are selected so that the temperature range covered contains the experimental T_m_ of the simulated protein. The three calculated ΔH_unf_ values are represented as a function of simulation temperature and the ΔCp_unf_ is calculated as the slope of a linear fit. For three-state proteins (e.g. apoFld), ΔCp_unf(F-to-I)_ and ΔCp_unf(I-to-U)_ are obtained as the temperature dependence of the calculated enthalpy changes of each unfolding transition.

The calculation of the protein stability curves (ΔG_unf_ as a function of temperature) is done through the Gibbs-Helmholtz equation^27^ (**Eq. 1**):

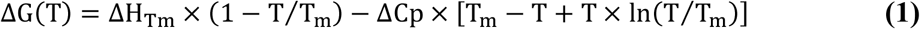

introducing the calculated ΔH_unf_ and ΔCp_unf_ values, and the reported experimental T_m_.

### MD-based specific workflow for calculation of unfolding energetics (ΔH_unf_, ΔCp_unf_ and ΔG_unf_) in holoproteins

In the case of holoproteins (non-covalent complexes of apoprotein and cofactor; e.g. holoFld), the ensemble-averaged enthalpy of the folded (bound) state, ⟨H_holo(F)_⟩, is obtained from simulations (40 replicas) each consisting of one molecule of holoFld solvated with water molecules and ions, as needed (**Figure 1** and **Figure 4a**). Similarly, the energetics of the unfolded (unbound) state is modeled from simulations (100 replicates) in which one unfolded protein molecule generated with ProtSA and one cofactor molecule (placed at a minimum distance of 3 nm from the protein) are put together in a box, where they are solvated in the same way (**Figure 4a**). The ensemble-averaged enthalpy of such boxes, ⟨H_apo(U)+cofactor_⟩, is obtained following the averaging scheme of the general workflow. Then, the unfolding enthalpy change is calculated as: ΔH_unf_ = ⟨H_apo(U)+cofactor_⟩ − ⟨H_holo(F)_⟩.

As required for this enthalpy change calculation by difference, the number of water molecules and ions in the box containing unfolded protein and cofactor must equate those in the box containing folded holoprotein (**SI Table 2**). The simulations are also performed at three different temperatures so that the unfolding ΔCp_unf_ can be obtained as the slope of a ΔH_unf_ versus temperature plot (**Figure 4a-b**).

Similarly to the case of apoproteins, the ΔH_unf_ and ΔCp_unf_ values calculated for the holoprotein can be combined with the experimental T_m_ to obtain the protein stability curves (ΔG_unf_ as a function of temperature). However, as the conformational stability of holoproteins is cofactor concentration dependent, a modified Gibbs-Helmholtz equation that takes into account the binding energetics (**SI Eq. 5**, see details in **SI Methods**) has been used to calculate the conformational stability as a function of temperature and concentration of free cofactor.

### Solvation conditions and MD simulation general details

Solvation conditions on the simulated proteins (i.e. protonation states and the number of ions added) were selected in each case to reproduce the experimental pH and ionic strength (IS) under which the experimental thermodynamics measurements were performed (see detailed information in **SI Methods** and **SI Table 2)**. Box dimensions were adopted from the diameter of the most elongated structure in the unfolded ensemble sampled for the protein in question plus a minimum distance of 1 nm from protein atoms to the simulation box edges. The MD simulation setup is similar to that described in Galano-Frutos et.al.^25^ (details are also given in **SI Table 1**). All the systems are simulated with the force field Charmm22 with CMAP correction (version 2.0)^18^ and the explicit water model Tip3p^28^: the most accurate force field/water model combination reported in previous work^25^. The Amber99SB-ILDN^17^ force field is again tested combined with Tip3p^28^ by modeling the apoFld unfolding thermodynamics. MD simulations are run and analyzed with Gromacs 2020 package^94^. The usage of short 2-ns productive trajectories in the workflow^25^ avoids the known issue of structure overcompaction in long simulations^16,25^ for force fields like Charmm22-CMAP^18^ and Amber99SB-ILDN^17^. In addition, the simulations performed here have been tested for overcompaction through analysis of the radius of gyration (Rg) evolution along the trajectories (**SI Table 3**). Results of this analysis confirm that no significant protein compaction takes place over the trajectories of the proteins simulated (**SI Table 3)**. The mutant variants here tested (of CI2 and lysozyme) have been modeled by replacing the wild-type residue by the new one using the mutator tool of Chimera (v.1.15)^95^, as no solved structures were available. No clashes were observed in the final mutant structures of lowest energy obtained after accommodating the new residues, which were taken as the starting structures in simulations of their folded states. In the case of the apoFld intermediate state, the representative model used (see below) was mutated back to its wild type sequence (Chimera v.1.15)^95^ in order to keep the same amino acids sequence as that of the other structural models used in simulations of apo and holoFld. No clashes were observed after this replacement either. Crystal waters and any other non-protein molecule were removed from the PDB structural models chosen (see below). Computational core facilities used in the current implementation consisted of a CPU-based cluster (170 Intel Xeon E5-2680v3 CPUs arranged in 85 nodes, https://cesar.unizar.es/hpc/), which was exploited through an Open MPI parallel framework (most simulations were launched on 32 cores: 32 OpenMP threads/1 MPI task).

### Structure models (PDBs) and coverage

The starting structures used to simulate the folded state of the proteins analyzed are those with the highest resolution available in the RCSB Protein Data Bank^92,93^ at the time of writing this manuscript. Namely: 1A2P (1.5 Å resolution)^96^ for barnase, 2SNS (1.5 Å)^97^ for nuclease (C-ter fragment), 2CI2 (2.0 Å)^98^ for CI2 (truncated form), 6LZM (1.8 Å)^75^ for lysozyme, 1L63 (1.75 Å)^99^ for pseudo-lysozyme, 1FTG (2.0 Å)^100^ for apoFld and 1FLV (2.0 Å)^101^ for holoFld. On the other hand, the thermal unfolding intermediate state of apoFld is represented by 2KQU^84^, a 20-model NMR ensemble of the Phe99Asn mutant previously shown to be a reliable representation of this state^83,84,102^ According to the reference sequences in UniProt^103^ the structural coverage of the solved sequences is 3-110 (barnase), 83-231 (nuclease C-terminal fragment), 20-84 (WT CI2 and Ile76Ala mutant), 1-162 (WT lysozyme and Ile3Glu mutant), 1-162 (pseudo-WT lysozyme), 3-170 (apo and holoFld)..

### FMN parameterization

Three different parameterizations of the FMN molecule (charge –2) have been tested. Namely, ‘Par.-1’ was done *ad hoc* assisted by the AmberTools20 package^104^ and the Gaussian 09 program^105^; ‘Par.-2’ was that reported by Schulten et.al.^106^; and ‘Par.-3’ was obtained through the SwissParam server^107^. FMN coordinates were extracted from the crystal structure of holoFld (PDB ID: 1FLV)^101^. For *ad-hoc* ‘Par.-1’, partial atomic charges were modeled with Gaussian 09 (HF/6-31G*), which were then fitted through the RESP method^108,109^ (with Antechamber)^104,110^, and finally, parameters obtained from the General Amber Force Field (GAFF^111^, Antechamber^104,110^). FMN coordinates were uploaded to SwissParam^107^ (‘Par.-3’) in mol2 format after adding hydrogen atoms. Except for van der Waals parameters, which are taken from the closest atom type in Charmm2, parameters and charges with this server are derived/taken from Merck Molecular Force Field (MMFF)^107^.

### Increased sampling for higher precision

As previously shown^25^, individual enthalpies (H^i^_F_, H^i^_U_ or H^i^_I_) of the simulated systems (i.e. boxes containing one protein molecule, several ions and thousands of water molecules) can mount to 10^5^ kcal/mol (negative values) or even higher (see **SI Table 4**). These big figures owe to the large number of water molecules present in the large simulation boxes required to solvate the unfolded conformations. In general, the larger the protein, the larger the negative enthalpy of the simulated box. Therefore, calculation of unfolding thermodynamics by difference requires a high precision (a low standard error in the calculation) to be able to assess the accuracy of the approach (the difference between experimental and *in silico* results). Since the enthalpy change of a partial thermal unfolding step of a protein (e.g. the apoFld F-to-I or I-to-U transitions) can be significantly lower than the enthalpy changes previously modeled for barnase and nuclease^25^ (see **Table 1**), a higher precision (standard error ≤ 10 kcal/mol) than that achieved in previous work^25^ is here guaranteed *a priori* by running a higher number of replicas (extended sampling, **Table 2**). For each system the minimum sample size necessary to meet such precision was estimated as reported (see the **Supplementary Information** of Galano-Frutos et al.^25^).

## Extended Data Figures

**Extended Data Figure 1.**
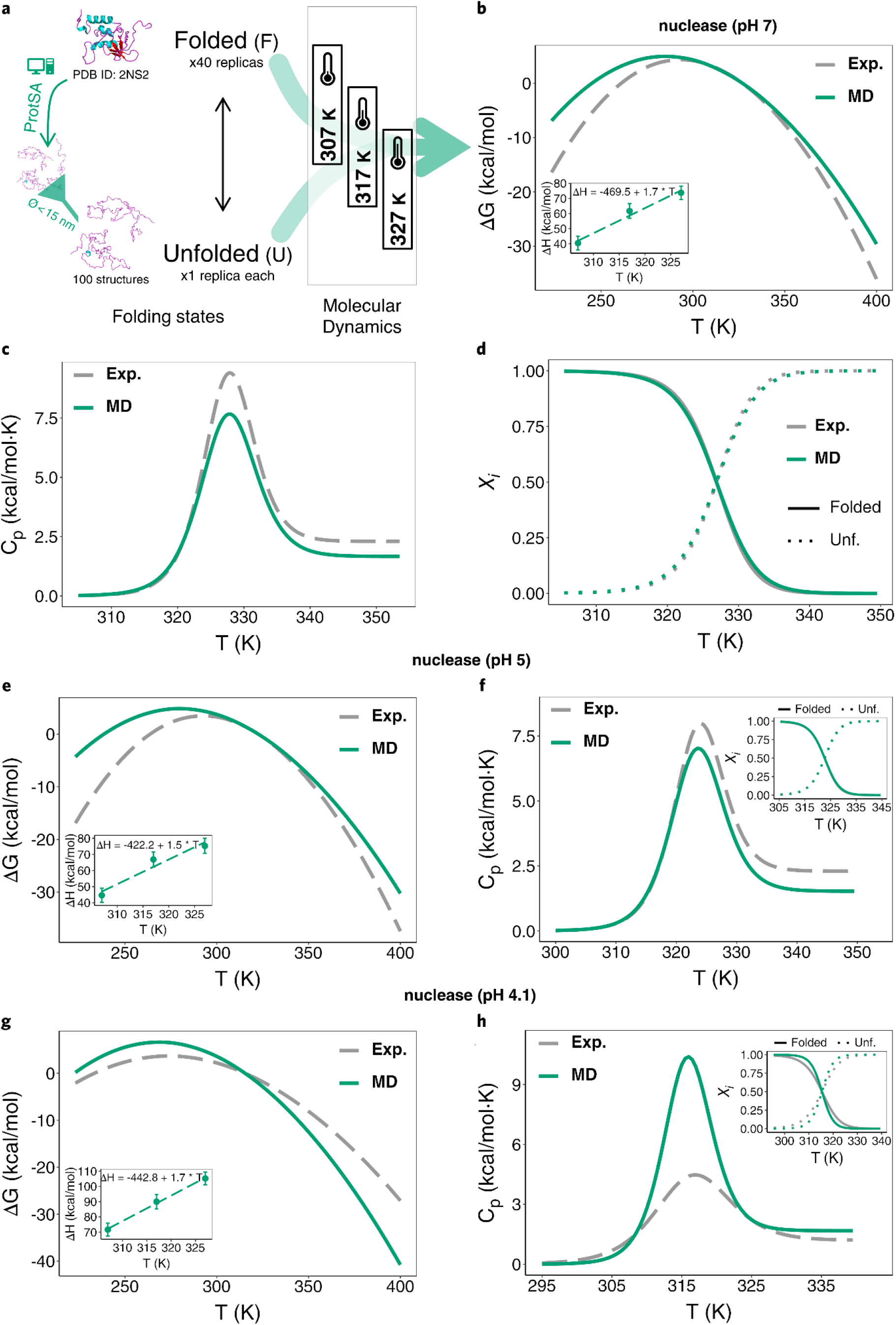
Simplified MD-based scheme and comparison with experimental results for nuclease. **a)** Protein models and folding states, number of structures (unfolded) and replicas (folded) simulated, diameter cutoff used to filter too-elongated unfolded structures obtained from ProtSA^9^ (left), and temperatures selected for the MD-based calculation (Charmm22-CMAP) of nuclease thermodynamics at pH 7.0, 5.0 and 4.1. **b-d)** Temperature profiles of ΔG_unf_, Cp_unf_ and *χ*_i_ (computed vs. experimental), respectively, obtained for nuclease simulated at pH 7.0. **e** and **f)** Temperature profiles of ΔG_unf_ and Cp_unf_ (computed vs. experimental), respectively, obtained for nuclease simulated at pH 5.0. **g** and **h)** Temperature profiles of ΔG_unf_ and Cp_unf_ (computed vs. experimental), respectively, obtained for nuclease simulated at pH 4.1. The ΔH_unf_ *vs*. T linear plot for pH ∼5.0 (and ∼4.1) appears as an inset in **e** (and **g**). The temperature profile of *χ*_*i*_ for pH 5.0 (and 4.1) is included as an inset in **f** (and **h**). The color-coding is indicated in the legends of the panels.

**Extended Data Figure 2.**
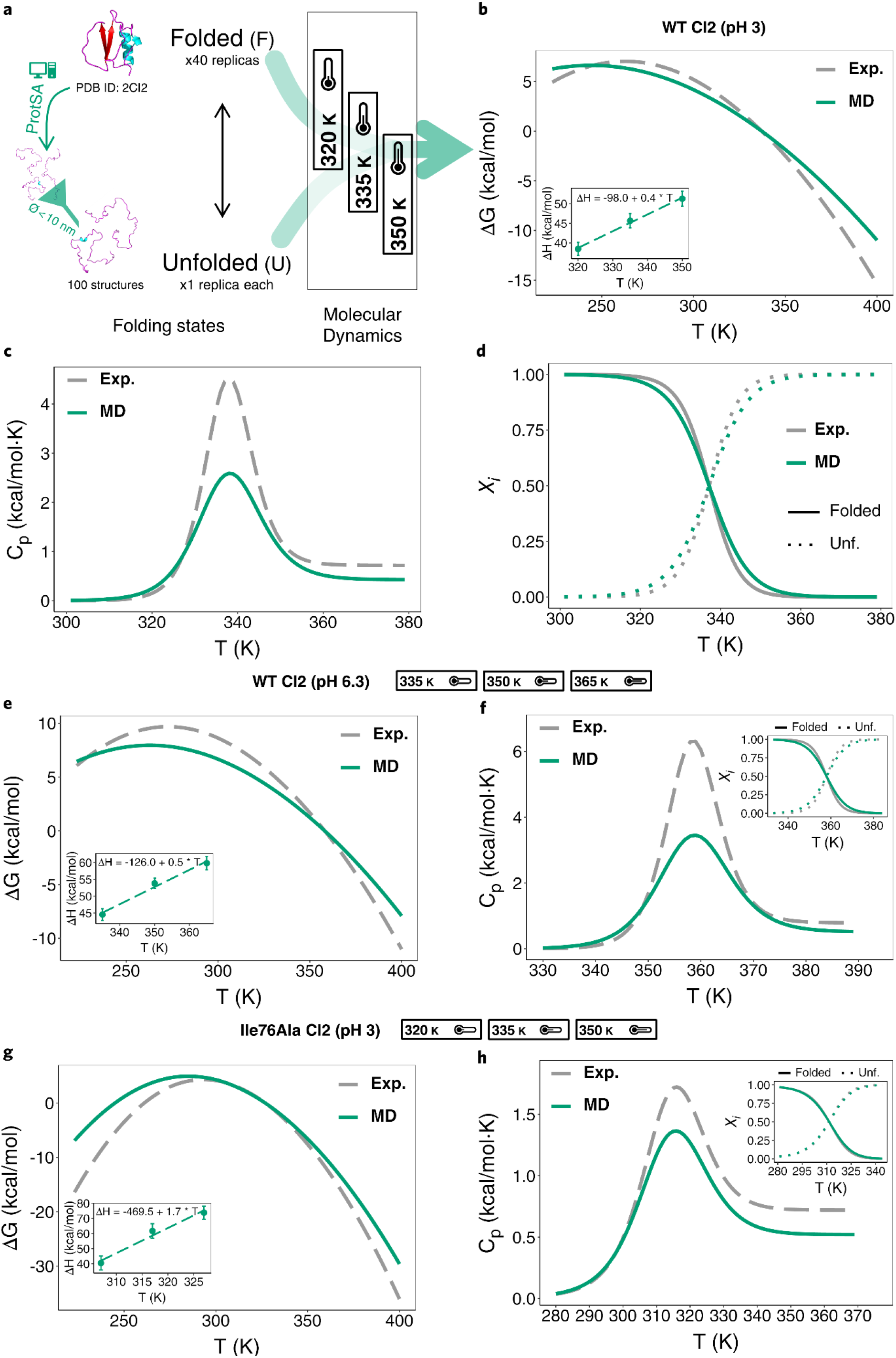
Simplified MD-based scheme and comparison with experimental results for CI2. **a)** Protein models and folding states, number of structures (unfolded) and replicas (folded) simulated, diameter cutoff used to filter too-elongated unfolded structures obtained from ProtSA^9^ (left), and temperatures selected for the MD-based calculation (Charmm22-CMAP) of CI2 thermodynamics. **b-d)** Temperature profiles of ΔG_unf_, Cp_unf_ and *χ*_i_ (*in silico* vs. experimental), respectively, obtained for WT CI2 at pH 3.0. Inset in **b** depicts the ΔH_unf_ vs. T linear plot with the fitted equation (the slope being ΔCp_unf_) obtained from the MD simulations. **e** and **f)** Temperature profiles of ΔG_unf_ and Cp_unf_ (computed vs. experimental), respectively, obtained for Ile76Ala at pH 3.0. The ΔH_unf_ vs. T linear plot appears as an inset in **e** and the temperature profile of *χ*_i_ is included as an inset in **f. g** and **h)** Temperature profiles of ΔG_unf_ and Cp_unf_ (computed vs. experimental), respectively, obtained for WT CI2 at pH 6.3. Insets in **g** and **h** similar to those in **e** and **f**, respectively. The color-coding is indicated in the legends of the panels.

**Extended Data Figure 3.**
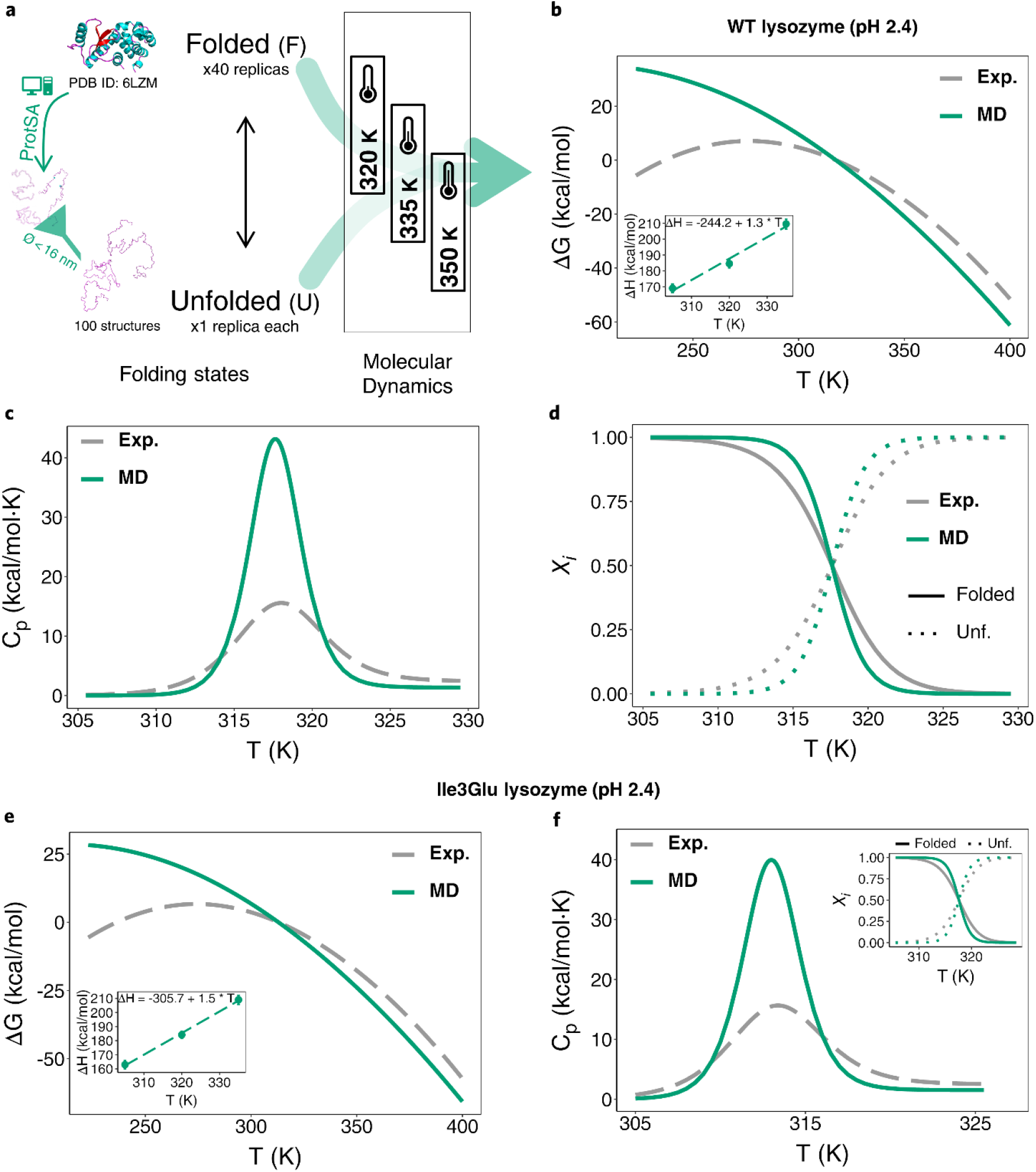
Simplified MD-based scheme and comparison with experimental results for lysozyme. **a)** The protein models and folding states, the number of structures (unfolded) and replicas (folded) simulated, the diameter cutoff used to filter too-elongated unfolded structures obtained from ProtSA^9^ (left), and temperatures selected for the MD-based calculation (Charmm22-CMAP) of thermodynamics of WT lysozyme and the variant Ile3Glu at pH 2.4. **b-d)** Temperature profiles of ΔG_unf_, Cp_unf_ and *χ*_*i*_ (*in silico* vs. experimental), respectively, obtained for WT lysozyme at pH 2.4. Inset in **b** depicts the ΔH_unf_ vs. T linear plot with the fitted equation (the slope being ΔCp_unf_) obtained from the MD simulations. **e** and **f)** Temperature profiles of ΔG_unf_ and Cp_unf_ (*in silico* vs. experimental), respectively, obtained for Ile3Glu at pH 2.4. Inset in **e** similar to that in **b**. The temperature profile of *χ*_*i*_ is included as an inset in **f**. The color-coding is indicated in the legends of the panels.

**Extended Data Figure 4.**
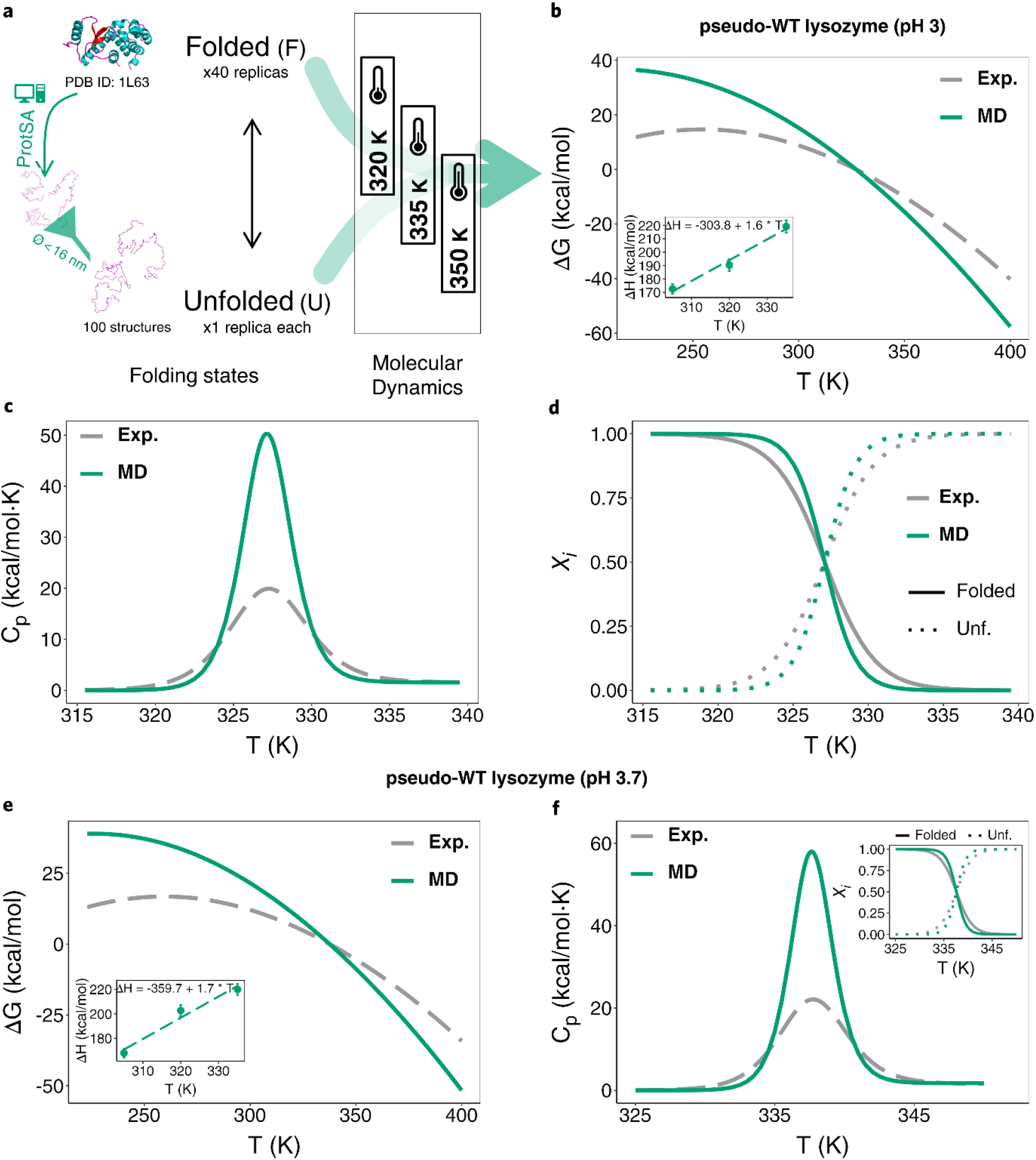
Simplified MD-based scheme and comparison with experimental results for pseudo-lysozyme. **a)** protein models and folding states, number of structures (unfolded) and replicas (folded) simulated, diameter cutoff used to filter too-elongated unfolded structures obtained from ProtSA^9^ (left), and temperatures selected for the MD-based calculation (Charmm22-CMAP) of pseudo-WT lysozyme thermodynamics. **b-d)** Temperature profiles of ΔG_unf_, Cp_unf_ and *χ*_*i*_ (computed vs. experimental), respectively, obtained for pseudo-WT lysozyme at pH 3.0. Inset in **b** depicts the ΔH_unf_ vs. T linear plot with the fitted equation (the slope being ΔCp_unf_) obtained from the MD simulations. **e** and **f)** Temperature profiles of ΔG_unf_ and Cp_unf_ (computed vs. experimental), respectively, obtained for pseudo-WT lysozyme at pH 3.7. Inset in **e** similar to that in **b**. The temperature profile of *χ*_*i*_ is included as an inset in **f**. The color-coding is indicated in the legends of the panels.

## ASSOCIATED CONTENT

### Supplementary Information

The Supplementary Information is available as a PDF document with the following sections:

#### SUPPLEMENTARY INFORMATION METHODS

Simulated Solvating Conditions.

Calculation of thermal stability (ΔGunf) of a three-state holoprotein (derivation of a Gibbs-Helmholtz-like equation).

#### SUPPLEMENTARY INFORMATION TABLES

Supplementary Information Tables 1-4.

#### SUPPLEMENTARY INFORMATION FIGURE

Supplementary Information Figure 1.

#### SUPPLEMENTARY INFORMATION REFERENCES

## Author Contributions

J.S. conceived and directed the investigation. J.J.G-F. and F.N-F. carried out and analysed the Molecular Dynamics simulations. J.J.G-F. and J.S. analysed data and wrote the manuscript.

## Notes

The authors declare no conflict of interest.

## Acknowledgements

We thank the Biocomputation and Complex Systems Physics Institute (BIFI) of the University of Zaragoza and the Red Española de Supercomputación (RES) for computing facilities granted to perform Molecular Dynamics simulations.

## Abbreviations

DSC: <u>D</u>ifferential <u>S</u>canning <u>C</u>alorimetry;
FMN: <u>F</u>lavin <u>M</u>ono<u>n</u>ucleotide;
IS: Ionic <u>S</u>trength;
LEM: <u>L</u>inear <u>E</u>xtrapolation <u>M</u>ethod;
MD: <u>M</u>olecular <u>D</u>ynamics;
NMR: <u>N</u>uclear <u>M</u>agnetic <u>R</u>esonance;
NPT: Isothermal-isobaric ensemble in MD simulations;
NVT: Canonical ensemble in MD simulations;
PBC: <u>P</u>eriodic <u>B</u>oundary <u>C</u>onditions;
PME: <u>P</u>article <u>M</u>esh <u>E</u>wald;
R_g_: Radius of gyration.

