## Supplementary Methods, Tables and Figure for "Calculation of Protein Folding Thermodynamics using Molecular Dynamics Simulations"

### Content


#### SUPPLEMENTARY INFORMATION METHODS

Simulated Solvating Conditions.

Calculation of thermal stability ($\Delta G_{unf}$) of a three-state holoprotein (derivation of a Gibbs-Helmholtz-like equation).

#### SUPPLEMENTARY INFORMATION TABLES

Supplementary Information Tables 1-4.

#### SUPPLEMENTARY INFORMATION FIGURE

Supplementary Information Figure 1.

#### SUPPLEMENTARY INFORMATION REFERENCES

### Supplementary Information Methods

**Simulated Solvating Conditions.** The protonation states set for the solution pH values simulated for the different proteins are shown in **SI Table 2**. Only protonation states for histidine, aspartic acid, glutamic acid, N-ter amine and C-ter carboxyl residues are included. The pKa values used to set the protonation states are only indicated in non-trivial cases. Both the indicated and non-shown pKa values were obtained from the PDB2PQR server[1,2] or, in the case of WT CI2[3], from NMR measurements. Given the pH values simulated, all basic residues (protonation and pKa values not shown in **SI Table 2**) were set to 'protonated'. Except for pseudo-lysozyme and nuclease (see footnote f in **Table 2**), salt concentration was set in the simulated systems to mimic the reported experimental ionic strength (see **Table 1** of the main text and **SI Table 2**). Those systems for which data reported under more than one experimental ionic strength were available, have been simulated either using one representative ionic strength value or the average. Ionic strengths have been modeled by adding the needed number of $Na^+$ and $Cl^-$ ions after setting the equivalent salt concentration through the Gromacs[4] tool *gmx genion*. **SI Table 2** also summarizes the protein charge, the number of water molecules and ions, and the box dimensions of the simulated systems. Importantly, solvating conditions for the unfolded ensembles (not shown in **SI Table 2**) are also adjusted to match the number of water molecules of their folded partner systems. Except for a small difference in the box dimensions, the solvating conditions for these unfolded ensembles are the same as those used for their folded counterparts (**SI Table 2**).

**Calculation of thermal stability ($\Delta G_{unf}$) of a three-state holoprotein (derivation of a Gibbs-Helmholtz-like equation).** According to the following reaction scheme:

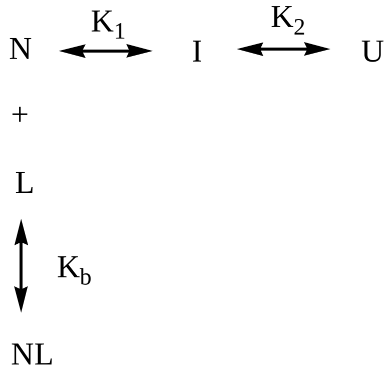


the unfolding equilibrium constant, K, of a holoprotein that binds a cofactor (or ligand) at a single binding site and whose apoprotein unfolds through a three-state mechanism (as it is the case of *Anabaena* holoFld) is:

$$K = \frac{[U]}{[N] + [NL]} = \frac{[U]}{[N]+K_b[N][L]} = \frac{[U]}{[N](1 + K_b[L])} = \frac{K_1K_2}{1 + K_b[L]}$$

where $K_1$ and $K_2$ are the equilibrium constants of the first (native to intermediate) and second (intermediate to unfolded) unfolding steps, respectively; $K_b$ is the binding constant of the complex formed between the native ($N$) protein and the cofactor or ligand ($L$), and the terms between brackets are the concentrations of free species in the equilibria.

Hence, the Gibbs free-energy change of the unfolding reaction can be written as:

$$\Delta G_{\text{holo(unf)}}(T) = -RT \cdot \ln(K_1K_2) + RT \cdot ln(1 + K_b(T) \cdot [L]) = \Delta G^0_{\text{apo(unf)}}(T) + RT \cdot ln(1 + K_b(T) \cdot [L]) = \Delta G^0_{(\text{Fol}\to\text{Int})}(T) + \Delta G^0_{(\text{Int}\to\text{Unf})}(T) + RT \cdot ln(1 + K_b(T) \cdot [L]) \quad \textbf{(1)},$$

where $\Delta G^0_{\text{apo(unf)}}(T)$ in the case of *Anabaena* Fld is the overall stability of the three-state apoprotein, and $\Delta G^0_{(\text{Fol}\to\text{Int})}(T)$ and $\Delta G^0_{(\text{Int}\to\text{Unf})}(T)$ are the Gibbs free-energies of the partial unfolding steps, which can be described through the Gibbs-Helmholtz equation (**Eq. 1** of the main text).

Wherever $K_b \gg 1/[L]$ **SI Eq. 1** can be approximated by:

$$\Delta G_{\text{holo(unf)}}(T) = \Delta G^0_{(\text{Fol}\to\text{Int})}(T) + \Delta G^0_{(\text{Int}\to\text{Unf})}(T) + RT \cdot ln(K_b(T) \cdot [L]) \quad \textbf{(2)}.$$

Binding constants are known to change with temperature[5] and the van't Hoff Equation[6] can be used to estimate their values from that at a reference temperature (e.g. 298.15 K):

$$ln\frac{K_b(T)}{K_b(298.15)} = \frac{-\Delta H_b + 298.15 \cdot \Delta Cp_b}{R} \cdot \left(\frac{1}{T} - \frac{1}{298.15}\right) + \frac{\Delta Cp_b}{R} ln\left(\frac{T}{298.15}\right) \quad \textbf{(3)}.$$

Combining **SI Eq. 2** and **3:**

$$\Delta G_{\text{holo(unf)}}(T) = \Delta G^0_{(\text{Fol}\to\text{Int})}(T) + \Delta G^0_{(\text{Int}\to\text{Unf})}(T) + RT \cdot \left\{lnK_b(298.15) + \frac{1}{R}\left[-\Delta H_b + 298.15 \cdot \Delta Cp_b \cdot \left(\frac{1}{T} - \frac{1}{298.15}\right) + \Delta Cp_b \cdot ln\left(\frac{T}{298.15}\right)\right]\right\} + RT \cdot ln[L] \quad \textbf{(4)},$$

which, at 1M of free ligand concentration becomes:

$$\Delta G_{\text{holo(unf)}}(T) = \Delta G^0_{(\text{Fol}\to\text{Int})}(T) + \Delta G^0_{(\text{Int}\to\text{Unf})}(T) + RT \cdot \left\{lnK_b(298.15) + \frac{1}{R}\left[-\Delta H_b + 298.15 \cdot \Delta Cp_b \cdot \left(\frac{1}{T} - \frac{1}{298.15}\right) + \Delta Cp_b \cdot ln\left(\frac{T}{298.15}\right)\right]\right\} \quad \textbf{(5)}.$$

**SI Eq. 5** provides the temperature dependence of the standard Gibbs free-energy change of the holoprotein. It should be indicated that the van't Hoff Equation cannot capture changes in the value of $K_b$ due to changes in the conformation of the native protein that may take place as the temperature increases. Therefore, we will assume here that **SI Eq. 5** may not hold above the first mid denaturation temperature of the apoprotein.

### Supplementary Information Tables

**Supplementary Information Table 1.** MD setup broken down by simulation steps[a]

| Simulation Step | General Settings (cutoffs) | PBC & Constraints | Step Setting | Thermodynamic Ensemble (baths) | Phys. Conditions | Simulated Time |
|---|---|---|---|---|---|---|
| Minimization | Integrator: steepest descent, Neighboring searching: grid, rcoul (PME): 1.0 nm, rvdw (cut-off): 1.0 nm | PBC: xyz, Constraints: none | Emtol: 1.0 kJ/mol | - | Temp: 0 K Press: 0 atm | max 20,000 steps |
| Volume adjustment[b] | Integrator: md, Neighboring searching: grid, rcoul (PME): 1.0 nm, rvdw (cut-off): 1.0 nm, vdw-modifier: potential-shift-verlet | PBC: xyz, Constraints: all-bonds | t-step: 1 fs | NPT Termost: v-rescale Barost: Berendsen | Temp: 0 K Press: 1 atm | 100 ps |
| Heating | | PBC: xyz, Constraints: all-bonds | t-step: 1 fs | NVT, Termost: Berendsen | Temp: Ini-T + ramp (6 x 50 K) Press: 0 atm | 7 x 50 ps |
| Equilibration 1 | | | t-step: 1 fs | NVT, Termost: v-rescale | Temp: Final-T Press: 0 atm | 150 ps |
| Equilibration 2 | | | t-step: 2 fs | NPT, Termost: v-rescale, Barost: Berendsen | Temp: Final-T Press: 1 atm | 250 ps |
| Equilibration 3 | | | | NPT, Termost: v-rescale, Barost: Parrinello-Rahman | | 250 ps |
| Production | | | | | | 2 ns |
| | | | | | | **Total:** 3.0 or 3.1 ns[c] |

[a] MD setup done with Gromacs package (v2020)[4].

[b] MD step designed to re-accommodate the water molecules inside the solvating box after having removed a few of them so that they match in number those in the reference system(s) (see footnote h in **SI Table 2**).

[c] Total simulated time along all the steps. 3.1 ns were simulated when the 'volume-adjustment' step was needed. 3.0 ns were simulated otherwise.

**Supplementary Information Table 2.** Solvating conditions and box dimensions in simulated systems[a]

| System[b] | pH | pKa(s)[c] | Protonation State[d] | Protein Charge[e] | Ioinic Strength[f] (mM) | No. Ions[g] | No. Waters[h] | Box Diameter[i] (nm) | Box Volume (nm³) |
|---|---|---|---|---|---|---|---|---|---|
| barnase | ~4.1 | Asp8:3.1 / Asp12:2.8 / Asp22:3.7 / Asp44:4.0 / Asp54:3.5 / Asp75:4.0 / Asp86:3.4 / Asp93:1.2 / Asp101:1.5 / Glu29:4.2 / Glu60:4.1 / Glu73:5.3 / His18:6.33 / His102:5.99 / N-ter:7.8 / C-ter:1.9 | Glu[prot] / His[prot] / Asp[unprot] / N-ter[prot] / C-ter[unprot] | +7 | 4.0 | 4 $Na^+$, 11 $Cl^-$ | 63,892 | 14 | 1,940 |
| nuclease[j] | 4.1 | Asp19:7.5 / Asp21:3.4 / Asp40:5.2 / Asp82:3.9 / Asp95:3.7 / Asp144:3.6 / Asp146:3.6 / Glu10:4.1 / Glu43:3.2 / Glu52:4.3 / Glu57:4.6 / Glu67:3.6 / Glu73:4.7 / Glu75:4.3 / Glu122:4.6 / Glu129:5.6 / Glu135:4.2 / Glu142:4.6 / His8:6.4 / His46:6.0 / His121:5.7 / His124:5.9 / N-ter:7.7 / C-ter:3.2 | Asp19[prot] / Asp21[unprot] / Asp40[prot] / Asp82[unprot] / Asp95[unprot] / Asp144[unprot] / Asp146[unprot] / Glu10[prot] / Glu43[unprot] / Glu52[prot] / Glu57[prot] / Glu67[unprot] / Glu73[prot] / Glu75[prot] / Glu122[prot] / Glu129[prot] / Glu135[prot] / Glu142[prot] / His8[prot] / His46[prot] / His121[prot] / His124[prot] / N-ter[prot] / C-ter[unprot] | +25 | neut. | 25 $Cl^-$ | 115,589 | 17 | 3,474 |
| | 5.0 | pKas: idem to nuclease pH 4.1 | Asp19[prot]/ His [prot] / N-ter[prot] / other Asp[unprot] / Glu[unprot] / C-ter[unprot] | +14 | neut. | 14 $Cl^-$ | 115,602 | 17 | 3,474 |
| | 7.0 | pKas: idem to nuclease pH 4.1 | Asp19[prot] / N-ter[prot] / other Asp[unprot] / Glu[unprot] / His[unprot]/ C-ter[unprot] | +11 | neut. | 11 $Cl^-$ | 115,602 | 17 | 3,474 |

**Supplementary Information Table 2.** Continuation…

| System[b] | pH | pKa(s)[c] | Protonation State[d] | Protein Charge[e] | Ioinic Strength[f] (mM) | No. Ions[g] | No. Waters[h] | Box Diameter[i] (nm) | Box Volume (nm³) |
|---|---|---|---|---|---|---|---|---|---|
| | 7.0 | pKas: idem to nuclease pH ~4.1 | Asp19[prot] / N-ter[prot] / other Asp[unprot] / Glu[unprot] / His[unprot]/ C-ter[unprot] | +11 | neut. | 11 $Cl^-$ | 115,602 | 17 | 3,474 |
| WT CI2[k] | 3.0 | Glu24*:2.9±0.1 / Glu27*:2.9±0.1 / Glu34*:3.5±0.1 / Glu35*:2.9±0.1 / Asp43*:2.4 ±0.4 / Glu46*:3.7±0.1 / Glu61*:3.1±0.0 / Asp65*:3.6±0.1 / Asp72*:2.5±0.1 / Glu79:4.8 / N-ter:7.9 / C-ter:1.4 | Glu24[unprot] / Glu27[unprot] / Glu34[prot] / Glu35[unprot] / Asp43[unprot] / Glu46[prot] / Glu61[prot] / Asp65[prot] / Asp72[unprot] / Glu79[prot] / N-ter[prot] / C-ter[unprot] | +5 | 1.9 | 1 $Na^+$, 6 $Cl^-$ | 30,950 | 11 | 941 |
| | 6.3 | N-ter:7.9 / C-ter:1.4 | Asp[unprot] / Glu[unprot] / N-ter[prot] / C-ter[unprot] | –1 | 33 | 19 $Na^+$, 18 $Cl^-$ | 30,902 | 11 | 941 |
| Ile76Ala CI2[k] | 3.0 | pKas: idem to WT CI2 pH 3.0 | Idem to WT CI2 pH 3.0 | +5 | 1.9 | 1 $Na^+$, 6 $Cl^-$ | 30,950 | 11 | 941 |
| WT lysozyme | 2.4 | Asp10:3.4 / Asp20:4.0 / Asp47:2.9 / Asp61:3.9 / Asp70:4.9 / Asp72:3.0 / Asp89:3.0 / Asp92:2.4 / Asp127:3.9 / Asp159:2.7 / Glu5:4.3 / Glu11:5.1 / Glu22:3.3 / Glu45:3.8 / Glu62:4.3 / Glu64:4.5 / Glu108:3.9 / Glu128:4.6 / His31:6.7 / N-ter:8.0 / C-ter:3.3 | Asp[prot] / Glu[prot] / His[prot] / N-ter[prot] / C-ter[prot] | +28 | 34 | 71 $Na^+$, 99 $Cl^-$ | 115,371 | 17 | 3,474 |

**Supplementary Information Table 2.** Continuation…

| System[b] | pH | pKa(s)[c] | Protonation State[d] | Protein Charge[e] | Ioinic Strength[f] (mM) | No. Ions[g] | No. Waters[h] | Box Diameter[i] (nm) | Box Volume (nm$^3$) |
|---|---|---|---|---|---|---|---|---|---|
| Ile3Glu lysozyme | 2.4 | Glu3:3.7 / Other pKas: idem to WT lysozyme pH 2.4 | Glu3$^{prot}$ / Other Glu$^{prot}$ / Asp$^{prot}$ / His$^{prot}$ / N-ter$^{prot}$ / C-ter$^{prot}$ | +28 | 34 | 71 $Na^+$, 99 $Cl^-$ | 115,371* | 17,01* | vol. adj |
| pseudo-WT lysozyme[k] | 3.0 | Asp10:3.4 / Asp20:3.9 / Asp47:2.7 / Asp61:4.0 / Asp70:4.8 / Asp72:3.2 / Asp89:2.8 / Asp92:2.1 / Asp127:3.9 / Asp159:2.6 / Glu5:4.4 / Glu11:5.0 / Glu22:3.3 / Glu45:3.7 / Glu62:2.8 / Glu64:4.7 / Glu108:3.9 / Glu128:4.6 / His31:6.6 / N-ter:8.0 / C-ter:3.3 | Asp10$^{prot}$ / Asp20$^{prot}$ / Asp47$^{unprot}$ / Asp61$^{prot}$ / Asp70$^{prot}$ / Asp72$^{prot}$ / Asp89$^{unprot}$ / Asp92$^{unprot}$ / Asp127$^{prot}$ / Asp159$^{unprot}$ / Glu5$^{prot}$ / Glu11$^{prot}$ / Glu22$^{prot}$ / Glu45$^{prot}$ / Glu62$^{unprot}$ / Glu64$^{prot}$ / Glu108prot / Glu128$^{prot}$ / His31$^{prot}$ / N-ter$^{prot}$ / C-ter$^{prot}$ | +23 | neut. | 23 $Cl^-$ | 115,531 | 17 | 3,474 |
| | 3.7 | pKas: idem to pseudo-WT lysozyme pH 3.0 | Asp10$^{unprot}$ / Asp72$^{unprot}$ / Glu22$^{unprot}$ / N-ter$^{prot}$ / C-ter$^{unprot}$ / other residues: idem to pseudo-WT lysozyme pH ~3.0 | +19 | neut. | 19 $Cl^-$ | 115,535 | 17 | 3,474 |

**Supplementary Information Table 2.** Continuation…

| System[b] | pH | pKa(s)[c] | Protonation State[d] | Protein Charge[e] | Ioinic Strength[f] (mM) | No. Ions[g] | No. Waters[h] | Box Diameter[i] (nm) | Box Volume ($nm^3$) |
|---|---|---|---|---|---|---|---|---|---|
| apoFld[†] | 7.0 | His35:5.4 / N-ter:7.2 / C-ter:3.4 | His35[unprot] / Asp[unprot] / Glu[unprot] / N-ter[prot] / C-ter[unprot] | –17 | 17 | 51 $Na^+$, 34 $Cl^-$ | 136,732 | 18 | 4,124 |
| apoFld[†] (intermediate) | 7.0 | His35:5.5 / N-ter:7.7 / C-ter:3.3 | His35[unprot] / Asp[unprot] / Glu[unprot] / N-ter[prot] / C-ter[unprot] | –17 | 17 | 51 $Na^+$, 34 $Cl^-$ | 136,732* | 19* | vol. adj |
| holoFld | 7.0 | His35:5.3 / N-ter:7.3 / C-ter:3.3 | His35[unprot] / Asp[unprot] / Glu[unprot] / N-ter[prot] / C-ter[unprot] | –19 | 17 | 53 $Na^+$, 34 $Cl^-$ | 145,423 | 19 | 4,524 |
| apoFld + FMN | 7.0 | His35:5.4 / N-ter:7.2 / C-ter:3.4 | His35[unprot] / Asp[unprot] / Glu[unprot] / N-ter[prot] / C-ter[unprot] | –19 | 17 | 53 $Na^+$, 34 $Cl^-$ | 145,423* | 20* | vol. adj |

[a] Solvating conditions for systems simulated using Charmm22-CMAP and Tip3p. Same conditions are used to simulate folded and unfolded systems, and for systems simulated with Amber99SB-ILDN and Tip3p. **Note:** in order to have the same charge and, therefore, the same number of ions and water molecules in folded and unfolded systems, identical pKa values are assumed for the folded and the unfolded state even if it is not strictly the case.

[b] Proteins or cofactors (FMN) simulated in this work. (†): systems which have been additionally simulated with Amber99SB-ILDN.

[c] pKa values for the native protein (folded) either obtained from the PDB2PQR (PROPKA 3.0[7]) server[1,2] (no errors given) or reported by Tan *et al*.[3] (indicated with an asterisk (*) next to the residue name).

[d] Protonation states resulted from the simulated pH and pKa values (equal for folded and unfolded states, see the Note in footnote a).

[e] Protein charge after setting protonation.

[f] Ionic strength (IS) established in the systems (by adding $Na^+$ and $Cl^-$ ions) based on the values used in the corresponding experimental determination of the thermal unfolding parameters. For pseudo-lysozyme, the experimental IS (see **Table 1**) are lower than those obtained after neutralizing the system prior to simulation so no extra ions have been added after neutralization. For nuclease, the unfolding thermodynamics measurements have been described to be independent of IS[8] and the experimental ISs used in the reference experimental works vary in a large range, so no extra ions have been added after neutralization. Individual and averaged ISs used in the

experimental measurements along with the papers of reference are given in **Table 1** of the main text.

[g] Number of ions added by Gromacs *gmx genion* tool after setting the required salt concentration.

[h] Number of water molecules finally simulated in the systems. (*): systems where the number of water molecules was initially higher (when setting the box diameter) but it was reduced to match the number of waters in the corresponding reference system(s). **Example:** to calculate the unfolding energetics of holoFld, two systems were involved: 1) boxes with folded holoFld and 2) boxes with unfolded Fld (apo) plus FMN unbound (see **Methods** and **Figure 4a** in the manuscript). In this case, the number of water molecules in the systems with unfolded apoFld plus FMN is initially set bigger than the numbers of this molecule previously settled in the folded holoFld system (the reference here). Then, as required in this protocol (based on calculation by difference of thermodynamics of unfolding states) a few water molecules have to be removed to equate their number in the reference system. In these cases, a ‘volume-adjustment’ step (indicated as ‘vol. adj.’ in the ‘Box volume’ column) is introduced in the preparation phase before the heating step (**SI Table 1**).

[i] Truncated rhombic dodecahedron boxes. Dimensions defined in each case by the unfolded structure with maximum diameter among those in the filtered unfolded ensemble: a minimum distance of 1 nm was left between the farthest atom of the protein from the center and the box edges. (*): Initial diameter settled before removing a few water molecules and adjusting the box volume.

[j] 83-231 (renumbered 1-149) C-ter fragment of nuclease as in PDB ID 2SNS (see **Methods** in the main text).

[k] Truncated CI2 structures lacking the first 19 amino acid residues.

[l] Lysozyme variant wherein cysteine residues at positions 54 and 97 are replaced by a threonine and an alanine, respectively.

**Supplementary Information Table 3.** Radius of gyration (Rg) at selected times over the simulated 2ns-trajectories, and statistical comparison

| **Protein (pH)[a]** | **Force field / FMN-Parameterization[b]** | **State[c]** | **Tem[d] (K)** | **Rg[0e] (nm)** | **Rg[f] ($t_{0/4}$) (nm)** | **Rg[f] ($t_{1/4}$) (nm)** | **Rg[f] ($t_{2/4}$) (nm)** | **Rg[f] ($t_{3/4}$) (nm)** | **Rg[f] ($t_{4/4}$) (nm)** | **P-value[g]** |
|---|---|---|---|---|---|---|---|---|---|---|
| barnase | Charmm22-CMAP | folded | 295 | 1.351 | 1.366 | 1.368 | 1.371 | 1.372 | 1.371 | 0.136 |
| barnase | Charmm22-CMAP | folded | 315 | 1.351 | 1.371 | 1.370 | 1.373 | 1.370 | 1.373 | 0.683 |
| barnase | Charmm22-CMAP | folded | 335 | 1.351 | 1.370 | 1.370 | 1.370 | 1.368 | 1.368 | 0.903 |
| barnase | Charmm22-CMAP | unfolded | 295 | 2.655 | 2.627 | 2.584 | 2.613 | 2.599 | 2.599 | 0.999 |
| barnase | Charmm22-CMAP | unfolded | 315 | 2.655 | 2.658 | 2.620 | 2.646 | 2.571 | 2.533 | 0.922 |
| barnase | Charmm22-CMAP | unfolded | 335 | 2.655 | 2.631 | 2.608 | 2.570 | 2.449 | 2.397 | 0.517 |
| nuclease (pH 4.1) | Charmm22-CMAP | folded | 307 | 1.540 | 1.555 | 1.557 | 1.557 | 1.561 | 1.567 | 0.291 |
| nuclease (pH 4.1) | Charmm22-CMAP | folded | 317 | 1.540 | 1.560 | 1.563 | 1.564 | 1.570 | 1.569 | 0.339 |
| nuclease (pH 4.1) | Charmm22-CMAP | folded | 327 | 1.540 | 1.565 | 1.566 | 1.569 | 1.565 | 1.565 | 0.961 |
| nuclease (pH 4.1) | Charmm22-CMAP | unfolded | 307 | 3.379 | 3.445 | 3.509 | 3.497 | 3.511 | 3.499 | 0.998 |
| nuclease (pH 4.1) | Charmm22-CMAP | unfolded | 317 | 3.379 | 3.593 | 3.614 | 3.674 | 3.664 | 3.637 | 0.989 |
| nuclease (pH 4.1) | Charmm22-CMAP | unfolded | 327 | 3.379 | 3.530 | 3.638 | 3.594 | 3.631 | 3.695 | 0.936 |
| nuclease (pH 5.0) | Charmm22-CMAP | folded | 307 | 1.540 | 1.572 | 1.580 | 1.575 | 1.569 | 1.566 | 0.572 |
| nuclease (pH 5.0) | Charmm22-CMAP | folded | 317 | 1.540 | 1.566 | 1.564 | 1.566 | 1.560 | 1.555 | 0.521 |
| nuclease (pH 5.0) | Charmm22-CMAP | folded | 327 | 1.540 | 1.578 | 1.565 | 1.563 | 1.553 | 1.554 | 0.009 |
| nuclease (pH 5.0) | Charmm22-CMAP | unfolded | 307 | 3.286 | 3.171 | 3.133 | 3.088 | 3.085 | 3.124 | 0.984 |
| nuclease (pH 5.0) | Charmm22-CMAP | unfolded | 317 | 3.286 | 3.165 | 3.154 | 3.134 | 3.182 | 3.090 | 0.986 |
| nuclease (pH 5.0) | Charmm22-CMAP | unfolded | 327 | 3.286 | 3.123 | 3.150 | 3.122 | 3.028 | 2.937 | 0.695 |

**Supplementary Information Table 3.** Continuation…

| Protein (pH)[a] | Force field / FMN-parameterization[b] | State[c] | Tem[d] (K) | Rg[0e] (nm) | Rg[f] ($t_{0/4}$) (nm) | Rg[f] ($t_{1/4}$) (nm) | Rg[f] ($t_{2/4}$) (nm) | Rg[f] ($t_{3/4}$) (nm) | Rg[f] ($t_{4/4}$) (nm) | P-value[g] |
|---|---|---|---|---|---|---|---|---|---|---|
| nuclease (pH 7.0) | Charmm22-CMAP | folded | 307 | 1.540 | 1.555 | 1.548 | 1.553 | 1.556 | 1.550 | 0.575 |
| nuclease (pH 7.0) | Charmm22-CMAP | folded | 317 | 1.540 | 1.563 | 1.560 | 1.554 | 1.552 | 1.550 | 0.219 |
| nuclease (pH 7.0) | Charmm22-CMAP | folded | 327 | 1.540 | 1.561 | 1.558 | 1.563 | 1.557 | 1.552 | 0.615 |
| nuclease (pH 7.0) | Charmm22-CMAP | unfolded | 307 | 3.379 | 3.381 | 3.396 | 3.406 | 3.396 | 3.366 | 1.000 |
| nuclease (pH 7.0) | Charmm22-CMAP | unfolded | 317 | 3.379 | 3.487 | 3.450 | 3.392 | 3.382 | 3.346 | 0.944 |
| nuclease (pH 7.0) | Charmm22-CMAP | unfolded | 327 | 3.379 | 3.448 | 3.457 | 3.448 | 3.435 | 3.382 | 0.994 |
| WT CI2 (pH 3.0) | Charmm22-CMAP | folded | 320 | 1.129 | 1.138 | 1.139 | 1.137 | 1.137 | 1.140 | 0.311 |
| WT CI2 (pH 3.0) | Charmm22-CMAP | folded | 335 | 1.129 | 1.136 | 1.141 | 1.137 | 1.137 | 1.136 | 0.124 |
| WT CI2 (pH 3.0) | Charmm22-CMAP | folded | 350 | 1.129 | 1.142 | 1.140 | 1.138 | 1.137 | 1.137 | 0.206 |
| WT CI2 (pH 3.0) | Charmm22-CMAP | unfolded | 320 | 2.251 | 2.303 | 2.199 | 2.190 | 2.146 | 2.150 | 0.827 |
| WT CI2 (pH 3.0) | Charmm22-CMAP | unfolded | 335 | 2.251 | 2.100 | 2.031 | 1.964 | 1.910 | 1.924 | 0.521 |
| WT CI2 (pH 3.0) | Charmm22-CMAP | unfolded | 350 | 2.251 | 2.054 | 2.050 | 1.987 | 1.958 | 1.921 | 0.809 |
| WT CI2 (pH 6.3) | Charmm22-CMAP | folded | 335 | 1.129 | 1.134 | 1.134 | 1.131 | 1.130 | 1.130 | 0.072 |
| WT CI2 (pH 6.3) | Charmm22-CMAP | folded | 350 | 1.129 | 1.134 | 1.131 | 1.130 | 1.131 | 1.125 | 0.010 |
| WT CI2 (pH 6.3) | Charmm22-CMAP | folded | 365 | 1.129 | 1.130 | 1.131 | 1.129 | 1.133 | 1.130 | 0.461 |
| WT CI2 (pH 6.3) | Charmm22-CMAP | unfolded | 335 | 2.251 | 2.273 | 2.215 | 2.100 | 2.080 | 2.002 | 0.391 |
| WT CI2 (pH 6.3) | Charmm22-CMAP | unfolded | 350 | 2.251 | 2.049 | 1.900 | 1.818 | 1.802 | 1.806 | 0.358 |
| WT CI2 (pH 6.3) | Charmm22-CMAP | unfolded | 365 | 2.251 | 1.938 | 1.966 | 1.903 | 1.848 | 1.740 | 0.339 |

**Supplementary Information Table 3.** Continuation…

| **Protein (pH)**[a] | **Force field / FMN-parameterization**[b] | **State**[c] | **Tem**[d] **(K)** | **Rg**[0e] **(nm)** | **Rg**[f] **($t_{0/4}$) (nm)** | **Rg**[f] **($t_{1/4}$) (nm)** | **Rg**[f] **($t_{2/4}$) (nm)** | **Rg**[f] **($t_{3/4}$) (nm)** | **Rg**[f] **($t_{4/4}$) (nm)** | **P-value**[g] |
|---|---|---|---|---|---|---|---|---|---|---|
| Ile76-Ala CI2 (pH 3.0) | Charmm22-CMAP | folded | 320 | 1.131 | 1.136 | 1.136 | 1.136 | 1.135 | 1.134 | 0.858 |
| Ile76-Ala CI2 (pH 3.0) | Charmm22-CMAP | folded | 335 | 1.131 | 1.136 | 1.136 | 1.135 | 1.136 | 1.136 | 0.964 |
| Ile76-Ala CI2 (pH 3.0) | Charmm22-CMAP | folded | 350 | 1.131 | 1.139 | 1.138 | 1.135 | 1.137 | 1.136 | 0.542 |
| Ile76-Ala CI2 (pH 3.0) | Charmm22-CMAP | unfolded | 320 | 2.309 | 2.149 | 2.110 | 2.099 | 2.107 | 2.103 | 0.994 |
| Ile76-Ala CI2 (pH 3.0) | Charmm22-CMAP | unfolded | 335 | 2.309 | 2.099 | 2.052 | 1.980 | 1.897 | 1.829 | 0.217 |
| Ile76-Ala CI2 (pH 3.0) | Charmm22-CMAP | unfolded | 350 | 2.309 | 2.052 | 2.035 | 2.032 | 1.944 | 1.922 | 0.852 |
| WT lysozyme (pH 2.4) | Charmm22-CMAP | folded | 305 | 1.652 | 1.702 | 1.702 | 1.704 | 1.696 | 1.697 | 0.656 |
| WT lysozyme (pH 2.4) | Charmm22-CMAP | folded | 320 | 1.652 | 1.708 | 1.706 | 1.711 | 1.697 | 1.701 | 0.299 |
| WT lysozyme (pH 2.4) | Charmm22-CMAP | folded | 335 | 1.652 | 1.707 | 1.706 | 1.708 | 1.711 | 1.699 | 0.698 |
| WT lysozyme (pH 2.4) | Charmm22-CMAP | unfolded | 305 | 3.644 | 3.442 | 3.473 | 3.474 | 3.491 | 3.432 | 0.998 |
| WT lysozyme (pH 2.4) | Charmm22-CMAP | unfolded | 320 | 3.644 | 3.438 | 3.449 | 3.422 | 3.470 | 3.513 | 0.993 |
| WT lysozyme (pH 2.4) | Charmm22-CMAP | unfolded | 335 | 3.644 | 3.413 | 3.428 | 3.477 | 3.419 | 3.403 | 0.991 |
| Ile3Glu lysozyme (pH 2.4) | Charmm22-CMAP | folded | 305 | 1.652 | 1.700 | 1.702 | 1.710 | 1.706 | 1.703 | 0.563 |
| Ile3Glu lysozyme (pH 2.4) | Charmm22-CMAP | folded | 320 | 1.652 | 1.705 | 1.708 | 1.707 | 1.705 | 1.710 | 0.972 |
| Ile3Glu lysozyme (pH 2.4) | Charmm22-CMAP | folded | 335 | 1.652 | 1.707 | 1.703 | 1.700 | 1.706 | 1.704 | 0.940 |
| Ile3Glu lysozyme (pH 2.4) | Charmm22-CMAP | unfolded | 305 | 3.700 | 3.538 | 3.551 | 3.593 | 3.602 | 3.610 | 0.994 |
| Ile3Glu lysozyme (pH 2.4) | Charmm22-CMAP | unfolded | 320 | 3.700 | 3.263 | 3.195 | 3.277 | 3.324 | 3.369 | 0.917 |
| Ile3Glu lysozyme (pH 2.4) | Charmm22-CMAP | unfolded | 335 | 3.700 | 3.310 | 3.264 | 3.323 | 3.315 | 3.281 | 0.998 |

**Supplementary Information Table 3.** Continuation…

| **Protein (pH)[a]** | **Force field / FMN-parameterization[b]** | **State[c]** | **Tem[d] (K)** | **$Rg^{0e}$ (nm)** | **$Rg^f$ ($t_{0/4}$) (nm)** | **$Rg^f$ ($t_{1/4}$) (nm)** | **$Rg^f$ ($t_{2/4}$) (nm)** | **$Rg^f$ ($t_{3/4}$) (nm)** | **$Rg^f$ ($t_{4/4}$) (nm)** | **P-value[g]** |
|---|---|---|---|---|---|---|---|---|---|---|
| pseudo-WT lysozyme (pH 3.0) | Charmm22-CMAP | folded | 305 | 1.651 | 1.707 | 1.706 | 1.704 | 1.703 | 1.707 | 0.987 |
| pseudo-WT lysozyme (pH 3.0) | Charmm22-CMAP | folded | 320 | 1.651 | 1.704 | 1.710 | 1.709 | 1.717 | 1.722 | 0.253 |
| pseudo-WT lysozyme (pH 3.0) | Charmm22-CMAP | folded | 335 | 1.651 | 1.711 | 1.709 | 1.717 | 1.711 | 1.714 | 0.963 |
| pseudo-WT lysozyme (pH 3.0) | Charmm22-CMAP | unfolded | 305 | 3.602 | 3.351 | 3.356 | 3.416 | 3.440 | 3.449 | 0.974 |
| pseudo-WT lysozyme (pH 3.0) | Charmm22-CMAP | unfolded | 320 | 3.602 | 3.317 | 3.316 | 3.319 | 3.305 | 3.301 | 1.000 |
| pseudo-WT lysozyme (pH 3.0) | Charmm22-CMAP | unfolded | 335 | 3.602 | 3.345 | 3.341 | 3.318 | 3.333 | 3.326 | 1.000 |
| pseudo-WT lysozyme (pH 3.7) | Charmm22-CMAP | folded | 305 | 1.651 | 1.665 | 1.671 | 1.665 | 1.664 | 1.666 | 0.897 |
| pseudo-WT lysozyme (pH 3.7) | Charmm22-CMAP | folded | 320 | 1.651 | 1.668 | 1.663 | 1.665 | 1.671 | 1.672 | 0.718 |
| pseudo-WT lysozyme (pH 3.7) | Charmm22-CMAP | folded | 335 | 1.651 | 1.662 | 1.672 | 1.674 | 1.681 | 1.680 | 0.099 |
| pseudo-WT lysozyme (pH 3.7) | Charmm22-CMAP | unfolded | 305 | 3.602 | 3.341 | 3.285 | 3.252 | 3.285 | 3.328 | 0.988 |
| pseudo-WT lysozyme (pH 3.7) | Charmm22-CMAP | unfolded | 320 | 3.602 | 3.259 | 3.253 | 3.240 | 3.305 | 3.324 | 0.983 |
| pseudo-WT lysozyme (pH 3.7) | Charmm22-CMAP | unfolded | 335 | 3.602 | 3.299 | 3.298 | 3.289 | 3.234 | 3.227 | 0.989 |
| apoFld | Charmm22-CMAP | folded | 305 | 1.465 | 1.486 | 1.486 | 1.487 | 1.488 | 1.489 | 0.126 |
| apoFld | Charmm22-CMAP | folded | 320 | 1.465 | 1.488 | 1.490 | 1.491 | 1.491 | 1.490 | 0.168 |
| apoFld | Charmm22-CMAP | folded | 335 | 1.465 | 1.492 | 1.493 | 1.493 | 1.492 | 1.494 | 0.849 |
| apoFld | Charmm22-CMAP | intermediate | 305 | 1.956 | 1.863 | 1.858 | 1.849 | 1.840 | 1.824 | 0.712 |
| apoFld | Charmm22-CMAP | intermediate | 320 | 1.956 | 1.867 | 1.839 | 1.828 | 1.802 | 1.803 | 0.042[h] |
| apoFld | Charmm22-CMAP | intermediate | 335 | 1.956 | 1.835 | 1.783 | 1.764 | 1.743 | 1.743 | 0.000[h] |

**Supplementary Information Table 3.** Continuation…

| **Protein (pH)[a]** | **Force field / FMN-Parameterization[b]** | **State[c]** | **Tem[d] (K)** | **Rg[0e] (nm)** | **Rg[f] ($t_{0/4}$) (nm)** | **Rg[f] ($t_{1/4}$) (nm)** | **Rg[f] ($t_{2/4}$) (nm)** | **Rg[f] ($t_{3/4}$) (nm)** | **Rg[f] ($t_{4/4}$) (nm)** | **P-value[g]** |
|---|---|---|---|---|---|---|---|---|---|---|
| apoFld | Charmm22-CMAP | unfolded | 305 | 3.522 | 3.412 | 3.378 | 3.340 | 3.236 | 3.150 | 0.833 |
| apoFld | Charmm22-CMAP | unfolded | 320 | 3.522 | 3.365 | 3.332 | 3.315 | 3.265 | 3.099 | 0.712 |
| apoFld | Charmm22-CMAP | unfolded | 335 | 3.522 | 3.254 | 3.294 | 3.242 | 3.183 | 3.051 | 0.823 |
| apoFld | Amber99SB-ILDN | folded | 305 | 1.465 | 1.481 | 1.478 | 1.480 | 1.479 | 1.481 | 0.443 |
| apoFld | Amber99SB-ILDN | folded | 320 | 1.465 | 1.482 | 1.481 | 1.480 | 1.481 | 1.482 | 0.854 |
| apoFld | Amber99SB-ILDN | folded | 335 | 1.465 | 1.480 | 1.482 | 1.481 | 1.481 | 1.482 | 0.941 |
| apoFld | Amber99SB-ILDN | intermediate | 305 | 1.956 | 1.879 | 1.858 | 1.843 | 1.842 | 1.832 | 0.266 |
| apoFld | Amber99SB-ILDN | intermediate | 320 | 1.956 | 1.852 | 1.849 | 1.856 | 1.837 | 1.834 | 0.875 |
| apoFld | Amber99SB-ILDN | intermediate | 335 | 1.956 | 1.869 | 1.839 | 1.835 | 1.815 | 1.806 | 0.050[h] |
| apoFld | Amber99SB-ILDN | unfolded | 305 | 3.526 | 3.559 | 3.544 | 3.588 | 3.578 | 3.623 | 0.999 |
| apoFld | Amber99SB-ILDN | unfolded | 320 | 3.526 | 3.276 | 3.256 | 3.244 | 3.240 | 3.232 | 1.000 |
| apoFld | Amber99SB-ILDN | unfolded | 335 | 3.526 | 3.290 | 3.275 | 3.290 | 3.332 | 3.281 | 0.999 |
| holoFld | Charmm22-CMAP/Par.-1 | folded | 305 | 1.460 | 1.485 | 1.485 | 1.485 | 1.484 | 1.486 | 0.817 |
| holoFld | Charmm22-CMAP/Par.-1 | folded | 320 | 1.460 | 1.485 | 1.486 | 1.486 | 1.485 | 1.488 | 0.361 |
| holoFld | Charmm22-CMAP/Par.-1 | folded | 335 | 1.460 | 1.490 | 1.489 | 1.489 | 1.489 | 1.489 | 0.930 |
| holoFld | Charmm22-CMAP/Par.-2 | folded | 305 | 0.457 | 0.461 | 0.460 | 0.459 | 0.460 | 0.459 | 0.222 |
| holoFld | Charmm22-CMAP/Par.-2 | folded | 320 | 0.457 | 0.461 | 0.460 | 0.460 | 0.460 | 0.461 | 0.924 |
| holoFld | Charmm22-CMAP/Par.-2 | folded | 335 | 0.457 | 0.459 | 0.460 | 0.460 | 0.459 | 0.460 | 0.833 |
| holoFld | Charmm22-CMAP/Par.-3 | folded | 305 | 1.460 | 1.486 | 1.484 | 1.485 | 1.483 | 1.485 | 0.742 |
| holoFld | Charmm22-CMAP/Par.-3 | folded | 320 | 1.460 | 1.486 | 1.486 | 1.485 | 1.488 | 1.484 | 0.312 |
| holoFld | Charmm22-CMAP/Par.-3 | folded | 335 | 1.460 | 1.488 | 1.486 | 1.488 | 1.486 | 1.489 | 0.592 |

[a] Protein or variant simulated. When a protein has been simulated at several pH values, the pH of the simulation is indicated between parentheses (see **SI Table 2**).

[b] Force field used in the simulation. In simulations of holoFld the FMN parameterization used is indicated (see section **Methods** of the main text).

[c] Folding state (folded, intermediate or unfolded) of the simulated systems.

[d] Temperature at which the trajectories where the Rg is analyzed were run.

[e] Radius of gyration of the crystal structure (see PDB IDs in the main text), average Rg of the initial structures selected from the unfolded ensembles generated by ProtSA[9], or average Rg of the NMR ensemble (PDB 2KQU[10]) representing the apoFld intermediate state.

[f] Average radius of gyration of the proteins (for all the simulated replicas per system) over time in the 2ns-production trajectories run, namely: $t_{0/4}$ at the beginning of the trajectory (t=0 ns), $t_{1/4}$ at the end of the first quartile of the trajectory (t=0.5 ns), $t_{2/4}$ at the middle of the trajectory (t=1.0 ns), $t_{3/4}$ at the end of the third quartile of the trajectory (t=1.5 ns), and $t_{4/4}$ at the end of the trajectory (t=2.0 ns).

[g] P-value of an ANOVA statistical test performed to compare the protein Rg means at the checkpoints indicated. Analysis done for assessing the Rg evolution over time in relation with the known issue of structure overcompaction reported[11,12] for force fields like Charmm22-CMAP[13].

[h] Significant P-values (for a 95 % of confidence) indicated. These values only appear in some systems of the apoFld intermediate state. In this state, the apoFld structure simulated (PDB 2KQU[10]) presents disordered regions –appearing quite stretched in the NMR models− that slightly shrink at the beginning of the simulation towards the central structured nucleus albeit without forming new secondary structures nor compacting the already folded structure of the protein.

**Supplementary Information Table 4.** Time-averaged enthalpies (⟨$H_{state}$⟩ ± S.E.) per system and simulated temperature. Values used in the protocol to calculate by difference the enthalpy change upon unfolding ($\Delta H_{unf}$)

| | | barnase kcal/mol (±SE) | nuclease kcal/mol (±SE) | | | CI2 kcal/mol (±SE) | | |
|---|---|---|---|---|---|---|---|---|
| **State** | **Temp[a]** | **WT pH ~4.1** | **WT pH 4.1** | **WT pH 5.0** | **WT pH 7.0** | **WT pH 3.0** | **WT pH 6.3** | **Ile76Ala pH 3.0** |
| Folded | T1 | −506,461 (±1) | −887,731 (±3) | −887,492 (±2) | −887,120 (±3) | −230,686 (±1) | −226,942 (±1) | −230,697 (±1) |
| | T2 | −482,056 (±1) | −865,762 (±4) | −865,523 (±2) | −865,151 (±4) | −221,874 (±1) | −218,164 (±1) | −221,885 (±1) |
| | T3 | −457,835 (±2) | −843,875 (±3) | −843,630 (±2) | −843,259 (±3) | −213,081 (±1) | −209,383 (±1) | −213,096 (±1) |
| Unfolded | T1 | −506,381 (±1) | −887,659 (±1) | −887,441 (±1) | −887,079 (±2) | −230,648 (±1) | −226,897 (±1) | −230,665 (±1) |
| | T2 | −481,951 (±1) | −865,672 (±1) | −865,452 (±2) | −865,090 (±1) | −221,828 (±1) | −218,111 (±1) | −221,845 (±1) |
| | T3 | −457,714 (±1) | −843,769 (±1) | −843,544 (±2) | −843,185 (±1) | −213,030 (±1) | −209,324 (±1) | −213,048 (±1) |

**Supplementary Information Table 4.** Continuation…

| | | lysozyme kcal/mol (±SE) | | pseudo-lysozyme kcal/mol (±SE) | |
|---|---|---|---|---|---|
| **State** | **Temp[a]** | **WT pH 2.4** | **Ile3Glu pH 2.4** | **WT* pH 3.0** | **WT* pH 3.7** |
| Folded | T1 | −906,054 (±2) | −906,103 (±2) | −893,020 (±2) | −892,948 (±1) |
| | T2 | −873,131 (±1) | −873,186 (±1) | −860,068 (±2) | −860,007 (±2) |
| | T3 | −840,396 (±2) | −840,452 (±2) | −827,300 (±2) | −827,228 (±2) |
| Unfolded | T1 | −905,884 (±1) | −905,940 (±1) | −892,847 (±1) | −892,780 (±2) |
| | T2 | −872,945 (±1) | −873,002 (±1) | −859,878 (±2) | −859,804 (±2) |
| | T3 | −840,186 (±1) | −840,243 (±1) | −827,081 (±2) | −827,008 (±1) |

**Supplementary Information Table 4.** Continuation…

| | | **apoFld kcal/mol (±SE)** | | **holoFld kcal/mol (±SE)** | | |
|---|---|---|---|---|---|---|
| **State** | **Temp**[a] | **Charmm22-CMAP** | **Amber99SB-ILDN** | **Charmm22-CMAP/Par.-1** | **Charmm22-CMAP/Par.-2** | **Charmm22-CMAP/Par.-3** |
| Folded | T1 | −1,055,434 (±4) | −1,055,875 (±3) | −1,122,685 (±3) | −1,122,628 (±3) | −1,122,702 (±4) |
| | T2 | −1,016,485 (±4) | −1,016,930 (±2) | −1,081,263 (±5) | −1,081,206 (±3) | −1,081,290 (±3) |
| | T3 | −977,741 (±4) | −978,187 (±4) | −1,040,057 (±5) | −1,039,997 (±3) | −1,040,081 (±4) |
| Unfolded | T1 | −1,055,390 (±1) | −1,055,849 (±2) | −1,122,677 (±3)[b] | −1,122,622 (±3)[b] | −1,122,692 (±3)[b] |
| | T2 | −1,016,406 (±1) | −1,016,867 (±2) | −1,081,214 (±4)[b] | −1,081,173 (±3)[b] | −1,081,239 (±3)[b] |
| | T3 | −977,622 (±1) | −0,978,086 (±2) | −1,039,958 (±4)[b] | −1,039,904 (±3)[b] | −1,039,994 (±3)[b] |
| Intermed iate | T1 | −1,055,414 (±2) | −1,055,840 (±2) | | | |
| | T2 | −1,016,445 (±2) | −1,016,877 (±2) | | | |
| | T3 | −977,676 (±2) | −978,111 (±2) | | | |

[a] Simulated temperatures for each system appear in **Table 2** of the main text. Since they are different for each simulated system, they are only listed here as T1, T2 and T3 for the sake of space.

[b] Refers to systems where the unfolded apoprotein and one unbound FMN molecule are simulated in the same box (see **Methods** and **Figure 4a** of the main text).

#### Supplementary Information Figure

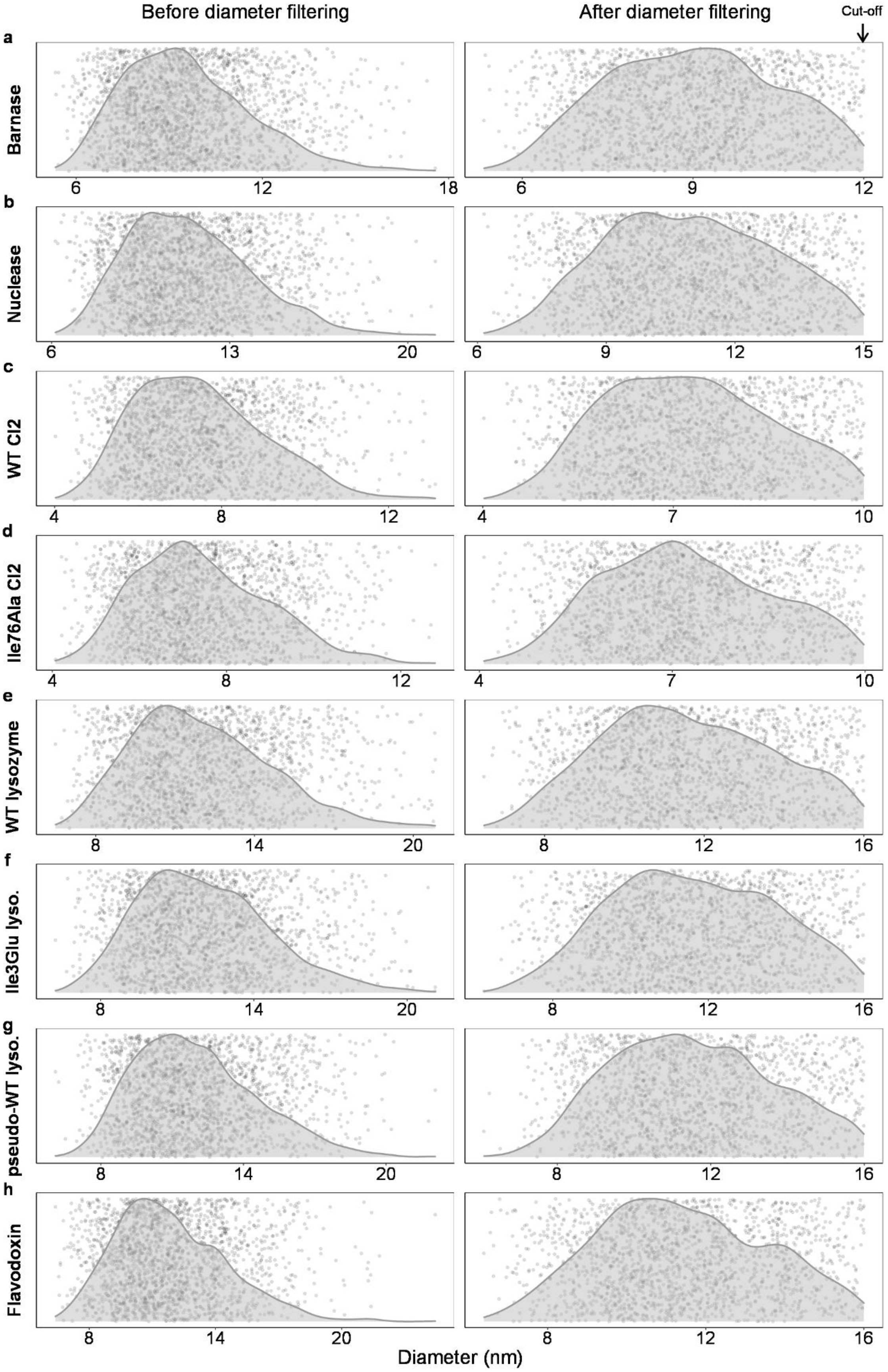

**Supplementary Information Figure 1.** Diameter distributions and cutoffs applied to filter out too elongated structures in the unfolded ensembles generated by ProtSA. Scatter and density plots obtained before (left) and after (right) diameter filtering for **a)** Barnase, **b)** Nuclease **c)** WT CI2, **d)** Ile76Ala CI2) **e)** WT lysozyme, **f)** Ile3Glu lysozyme, **g)** pseudo-WT lysozyme and **h)** apoFld. Maximum diameter (applied cutoff) kept for the unfolded ensembles are the maximum values in the depicted *x*-axis scale in plots at the right-hand side (indicated in the top right-hand corner of the figure).

#### Supplementary Information References


1. Dolinsky, T. J., Nielsen, J. E., McCammon, J. A. & Baker, N. A. PDB2PQR: An automated pipeline for the setup of Poisson-Boltzmann electrostatics calculations. *Nucleic Acids Res.* **32**, (2004).
2. Dolinsky, T. J. *et al.* PDB2PQR: Expanding and upgrading automated preparation of biomolecular structures for molecular simulations. *Nucleic Acids Res.* **35**, (2007).
3. Tan, Y. J., Oliveberg, M., Davis, B. & Fersht, A. R. Perturbed pKA-values in the denatured states of proteins. *J. Mol. Biol.* **254**, 980–992 (1995).
4. Van Der Spoel, D. *et al.* GROMACS: Fast, flexible, and free. *Journal of Computational Chemistry* vol. 26 1701–1718 (2005).
5. Clark, A. F., Hogg, R. W. & Gerken, T. A. Proton Nuclear Magnetic Resonance Spectroscopy and Ligand Binding Dynamics of the Escherichia coli l-Arabinose Binding Protein. *Biochemistry* **21**, 2227–2233 (1982).
6. Fukada, H., Sturtevant, J. M. & Quiocho, F. A. Thermodynamics of the binding of L-arabinose and of D-galactose to the L-arabinose-binding protein of Escherichia coli. *J. Biol. Chem.* **258**, 13193–13198 (1983).
7. Søndergaard, C. R., Olsson, M. H. M., Rostkowski, M. & Jensen, J. H. Improved treatment of ligands and coupling effects in empirical calculation and rationalization of pKa values. *J. Chem. Theory Comput.* **7**, 2284–2295 (2011).
8. Carra, J. H., Anderson, E. A. & Privalov, P. L. Thermodynamics of staphylococcal nuclease denaturation. I. The acid- denatured state. *Protein Sci.* **3**, 944–951 (1994).
9. Estrada, J., Bernadó, P., Blackledge, M. & Sancho, J. ProtSA: A web application for calculating sequence specific protein solvent accessibilities in the unfolded ensemble. *BMC Bioinformatics* **10**, 104 (2009).
10. Ayuso-Tejedor, S. *et al.* Design and Structure of an Equilibrium Protein Folding Intermediate:

A Hint into Dynamical Regions of Proteins. *J. Mol. Biol.* **400**, 922–934 (2010).

11. Galano-Frutos, J. J. & Sancho, J. Accurate Calculation of Barnase and SNase Folding Energetics Using Short Molecular Dynamics Simulations and an Atomistic Model of the Unfolded Ensemble: Evaluation of Force Fields and Water Models. *J. Chem. Inf. Model.* **59**, 4350–4360 (2019).

12. Robustelli, P., Piana, S. & Shaw, D. E. Developing a molecular dynamics force field for both folded and disordered protein states. *Proc. Natl. Acad. Sci. U. S. A.* **115**, E4758–E4766 (2018).

13. Mackerell, A. D., Feig, M. & Brooks, C. L. Extending the treatment of backbone energetics in protein force fields: Limitations of gas-phase quantum mechanics in reproducing protein conformational distributions in molecular dynamics simulation. *J. Comput. Chem.* **25**, 1400–1415 (2004).
